# Poor Quality Vβ Recombination Signal Sequences Enforce TCRβ Allelic Exclusion by Limiting the Frequency of Vβ Recombination

**DOI:** 10.1101/2020.01.20.913046

**Authors:** Glendon S. Wu, Katherine S. Yang-Iott, Morgann A. Reed, Katharina E. Hayer, Kyutae D. Lee, Craig H. Bassing

**Author notes:** Corresponding Author: Craig H. Bassing, Ph.D., Children’s Hospital of Philadelphia, 4054 Colket Translational Research Building, 3501 Civic Center Blvd., Philadelphia, PA 19104.

## Abstract

Monoallelic expression (allelic exclusion) of T and B lymphocyte antigen receptor genes is achieved by the assembly of a functional gene through V(D)J recombination on one allele and subsequent feedback inhibition of recombination on the other allele. There has been no validated mechanism for how only one allele of any antigen receptor locus assembles a functional gene prior to feedback inhibition. Here, we demonstrate that replacement of a single Vβ recombination signal sequence (RSS) with a better RSS increases Vβ rearrangement, reveals *Tcrb* alleles compete for utilization in the αβ T cell receptor (TCR) repertoire, and elevates the fraction of αβ T cells expressing TCRβ protein from both alleles. The data indicate that poor qualities of Vβ RSSs for recombination with Dβ and Jβ RSSs enforces allelic exclusion by stochastically limiting the incidence of functional Vβ rearrangements on both alleles before feedback inhibition terminates Vβ recombination.

## INTRODUCTION

Monoallelic gene expression is common, underlying genomic imprinting and X-chromosome activation in many cell types and tissue-specific allelic exclusion of olfactory neuron receptors and lymphocyte antigen receptors. Each of these programs has an initiation and a maintenance phase and involves epigenetic-based transcriptional silencing (Khamlichi and Feil, 2018). Lymphocyte antigen receptor (AgR) allelic exclusion involves additional levels of regulation due to obligate assembly of AgR genes through V(D)J recombination. In the germline, T cell receptor (TCR) and immunoglobulin (Ig) AgR loci are comprised of noncontiguous variable (V), joining (J), and, in some cases diversity (D), gene segments. Within developing T and B cells, the RAG1/RAG2 endonuclease cleaves at recombination signal sequences (RSSs) flanking V, D, and J segments to generate V(D)J rearrangements that assemble functional Ig and TCR genes (Bassing et al., 2002; Schatz and Swanson, 2011). Due to imprecision in repair of RAG DNA double strand breaks (DSBs), only about one-third of V(D)J rearrangements assembles an in-frame exon. In the absence of any regulation, the frequent assembly of out-of-frame rearrangements and requirement of AgR protein expression for T and B cell development dictates that biallelic expression of any TCR or Ig gene can occur in at most 20% of lymphocytes (Figure S1A) (Brady et al., 2010b; Mostoslavsky et al., 2004). However, TCRβ (*Tcrb*), IgH (*Igh*), and Igκ (*Igk*) loci exhibit more stringent allelic exclusion that is enforced by the assembly of a functional in-frame V(D)J rearrangement on one allele and subsequent feedback inhibition of V rearrangements on the other allele (Brady et al., 2010b; Levin-Klein and Bergman, 2014; Mostoslavsky et al., 2004; Outters et al., 2015; Vettermann and Schlissel, 2010). Thus, in ~60% of T or B cells only one V-to-(D)J rearrangement is found at each of these loci, while ~40% of T or B cells exhibit V-to-(D)J recombination on both alleles where typically only one rearrangement is in-frame (Figure S1B).

AgR gene assembly and expression are interdependently regulated with T and B cell development. CD4^-^CD8^-^ double-negative (DN) thymocytes and pro-B cells induce transcription, accessibility, and compaction of *Tcrb* or *Igh* loci, respectively (Brady et al., 2010b; Shih and Krangel, 2013). This accessibility allows RAG to bind at D and J segments, forming a focal recombination center (RC) in which D-to-J recombination occurs (Ji et al., 2010). Subsequently, a single V segment rearranges to a DJ complex on only one allele at a time (Brady et al., 2010b; Mostoslavsky et al., 2004; Outters et al., 2015; Vettermann and Schlissel, 2010). This V-to-DJ recombination step likely requires V segment accessibility and locus compaction to place V segments in spatial proximity with the RC (Brady et al., 2010b; Shih and Krangel, 2013). DSBs induced in DN thymocytes or pro-B cells repress RAG expression (Fisher et al., 2017), which may transiently inhibit further *Tcrb* and *Igh* recombination (Steinel et al., 2014). Cells that assemble an out-of-frame VDJ rearrangement on the first allele can attempt V recombination on the other allele (Brady et al., 2010b; Koralov et al., 2006; Lee and Bassing, 2020; Mostoslavsky et al., 2004; Outters et al., 2015; Vettermann and Schlissel, 2010). Following an in-frame VDJ rearrangement, resultant TCRβ or IgH proteins signal down-regulation of RAG expression and differentiation of CD4^+^CD8^+^ double-positive (DP) thymocytes or pre-B cells (von Boehmer and Melchers, 2010). These cells re-express RAG and recombine *Tcra* or *Igk* loci, but block further V-to-DJ rearrangements at *Tcrb* and *Igh* loci as a result of permanent feedback inhibition likely mediated through silencing of unrearranged V segments and locus de-contraction (Brady et al., 2010b; Majumder et al., 2015; Shih and Krangel, 2013). DP thymocytes assemble VJ rearrangements on both *Tcra* alleles until at least one allele yields a protein that forms an αβ TCR, which can signal differentiation of CD4^+^ or CD8^+^ single-positive (SP) thymocytes that are naïve mature αβ T cells (von Boehmer and Melchers, 2010). Pre-B cells assemble VJ rearrangements on one *Igk* allele at a time, and resulting RAG DSBs signal transient feedback inhibition of recombination on the other allele (Steinel et al., 2013). The formation and positive selection of an IgH/Igκ B cell receptor signals permanent feedback inhibition of V_κ_ recombination and maturation of κ^+^ B cells (von Boehmer and Melchers, 2010). As a result of these interdependent controls of lymphocyte development and V(D)J recombination between alleles, ~90% of αβ T cells and ~97% of κ^+^ B cells express only one type of AgR (Brady et al., 2010b).

While feedback inhibition mechanisms have been demonstrated experimentally, there have been no proven mechanisms for monoallelic assembly of a functional AgR gene prior to feedback inhibition. Both deterministic and stochastic models have been proposed to explain asynchronous timing of V-to-(D)J recombination between alleles of *Tcrb, Igh*, and *Igk* loci (Brady et al., 2010b; Levin-Klein and Bergman, 2014; Mostoslavsky et al., 2004; Outters et al., 2015; Vettermann and Schlissel, 2010). Deterministic models invoke that mechanisms predominantly activate one allele for V rearrangement and activate the second allele only if the first fails to assemble a functional gene. In contrast, stochastic models posit that both alleles are simultaneously active and mechanisms lower recombination efficiency, making it unlikely that both alleles assemble genes before feedback inhibition from one allele ceases V rearrangements. At least for *Igk*, asynchronous replication of homologous AgR alleles initiates in lymphoid progenitors, is clonally maintained, and correlates with preferential accessibility and recombination of the early replicating allele (Farago et al., 2012; Mostoslavsky et al., 2001). These findings suggest that asynchronous replication is a deterministic mechanism for monoallelic initiation of V recombination. In the lymphocyte lineage and developmental stage that *Tcrb, Igh*, or *Igk* loci recombine, their individual alleles frequently reside in different nuclear locations with V(D)J-rearranged alleles underrepresented at transcriptionally repressive nuclear structures (Chan et al., 2013; Hewitt et al., 2009; Schlimgen et al., 2008; Skok et al., 2007). The positioning of an allele at these structures by deterministic or stochastic means could block V rearrangements by suppressing accessibility, RAG binding, and/or locus compaction (Chan et al., 2013; Chen et al., 2018; Hewitt et al., 2009; Schlimgen et al., 2008; Skok et al., 2007). Sequence features conserved among Vβ and V_H_ RSSs, but not present in Dβ, J_H_, Vα, or V_κ_ RSSs, have been proposed to render Vβ and V_H_ recombination inefficient, thereby stochastically lowering the likelihood of near-simultaneous V rearrangements on both alleles (Liang et al., 2002). Although these proposed mechanisms may dictate monoallelic AgR gene assembly before enforcement of feedback inhibition, none have been validated by experimentally demonstrating causal relationships.

The mouse *Tcrb* locus offers a powerful physiological platform to elucidate potential contributions of RSSs in monoallelic assembly and expression of functional AgR genes. *Tcrb* has 23 functional Vβs located 250-735 kb upstream of the Dβ1-Jβ1-Cβ1 and Dβ2-Jβ2-Cβ2 clusters, each of which has one Dβ and six functional Jβs (Figure 1A) (Glusman et al., 2001; Malissen et al., 1986). The locus has another Vβ (*V31*) located 10 kb downstream of Cβ2 and in opposite transcriptional orientation from other *Tcrb* coding sequences (Glusman et al., 2001; Malissen et al., 1986). RSSs consist of a semi-conserved heptamer and nonamer separated by a generally non-conserved 12 or 23 nucleotide spacer (Schatz and Swanson, 2011). Upon binding an RSS, RAG adopts an asymmetric tilt conformation that ensures the capture of a second RSS of differing length and bends each RSS by inducing kinks in their spacers (Kim et al., 2018; Ru et al., 2015). *In vitro*, ~40% of synapses between RSSs with consensus heptamers and nonamers proceed to cleavage (Lovely et al., 2015), and natural variations of heptamers, spacers, and nonamers can have major effects on recombination levels (Akira et al., 1987; Connor et al., 1995; Gauss and Lieber, 1992; Hesse et al., 1989; Larijani et al., 1999; Livak et al., 2000; Nadel et al., 1998; Olaru et al., 2004; Ramsden and Wu, 1991; VanDyk et al., 1996; Wei and Lieber, 1993). The only *in vivo* confirmation that natural RSS variations influence recombination levels is in the *Tcrb* locus (Bassing et al., 2000; Horowitz and Bassing, 2014; Jung et al., 2003; Sleckman et al., 2000; Wu et al., 2003; Wu et al., 2007). Vβs are flanked by 23-RSSs, Jβs by 12-RSSs, and Dβs by 5’12-RSSs and 3’23-RSSs (Glusman et al., 2001). Direct Vβ-to-Jβ rearrangements are permitted by the 12/23 rule; however, they rarely occur due to the inherent inefficiency of recombination between Vβ and Jβ RSSs (Bassing et al., 2000; Jung et al., 2003; Tillman et al., 2003; Wu et al., 2003; Wu et al., 2007). The recombination strength of a *Tcrb* RSS is a property determined at the biochemical level by its interactions with a partner RSS, the RAG endonuclease, and HMGB1 proteins that bend DNA (Banerjee and Schatz, 2014; Drejer-Teel et al., 2007; Jung et al., 2003). In this context, 3’Dβ RSSs are at least 10-fold better than Vβ RSSs at recombining with 5’Dβ RSSs *in vitro* (Banerjee and Schatz, 2014; Drejer-Teel et al., 2007; Jung et al., 2003). Accordingly, replacement of an endogenous *V31* RSS with the 3’Dβ1 RSS increases the percentage of αβ T cells expressing V31^+^ TCRβ chains due to the elevated recombination level of *V31* relative to other Vβ segments (Horowitz and Bassing, 2014; Wu et al., 2003).

**Figure 1.**
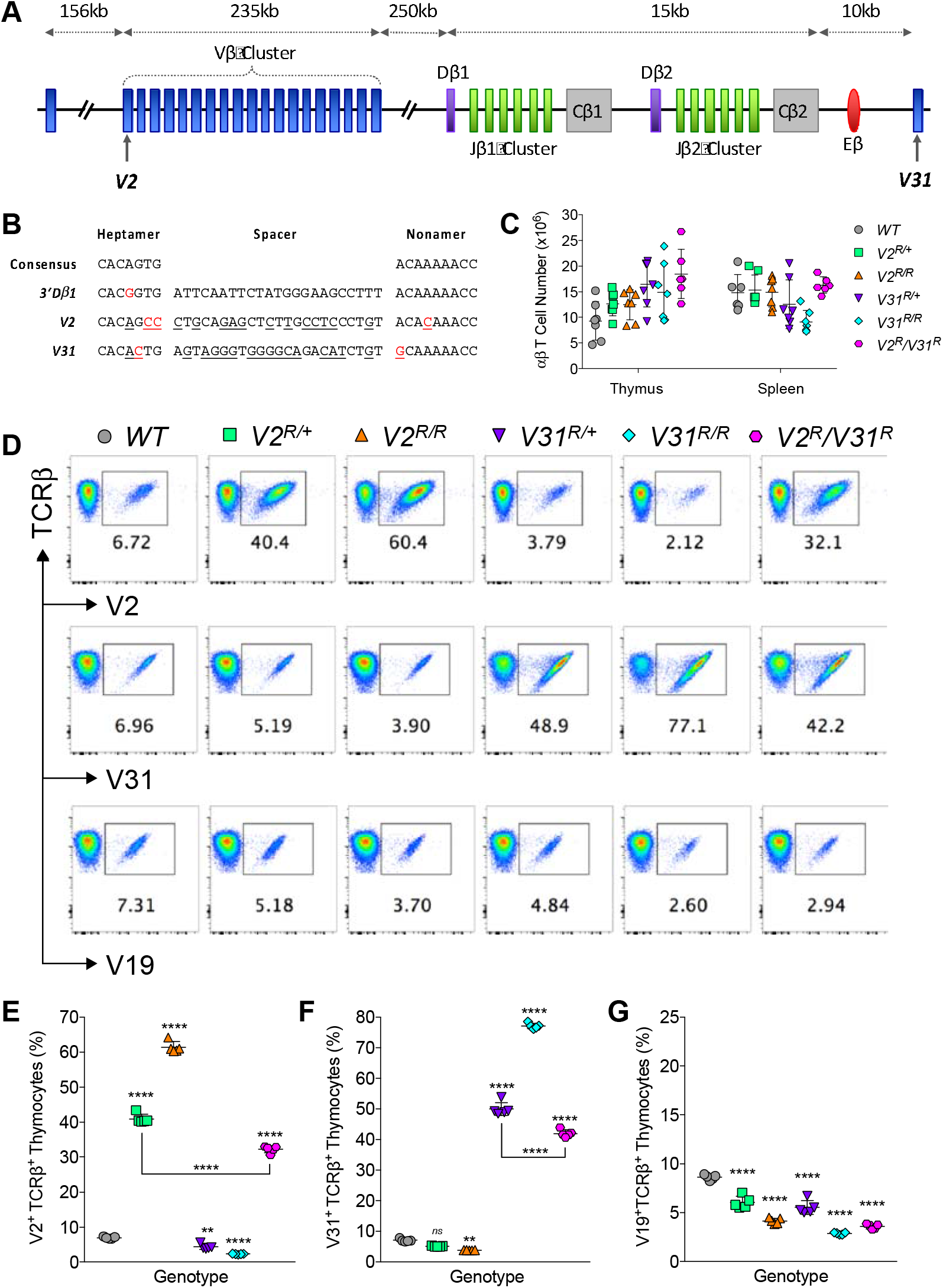
Increased Utilization of 3’Dβ1 RSS-replaced Vβ Segments on αβ T Cells. (A) Schematic of the *Tcrb* locus and relative positions of V, D, and J segments, C exons, and the Eβ enhancer. Not drawn to scale. (B) Sequences of a consensus heptamer and nonamer and the 3’Dβ1, V2, and V31 RSSs. Differences relative to the consensus heptamer and nonamer are indicated in red. Differences of each Vβ RSS relative to the 3’Dβ1 RSS are underlined. (C) Total numbers of SP thymocytes and splenic αβ T cells (n ≥ 6). (D) Representative plots of SP thymocytes expressing V2^+^, V31^+^, or V19^+^ TCRβ chains. (E-G) Quantification of V2^+^ (E), V31^+^ (F), or V19^+^ (G) SP thymocytes, refer to legend in (D). (n = 5, one-way ANOVA, multiple post-tests compared to *WT*, unless indicated by bars, and *p*-values are corrected for multiple tests. ns=not significant, \*\**p*<0.01, \*\*\*\**p*<0.0001). All quantification plots show mean ± SD.

To determine the potential roles of *Tcrb* RSSs in governing TCRβ allelic exclusion, we made and studied mice carrying replacement(s) of their endogenous *V31* and/or *Trbv2* (*V2*) RSSs with a better 3’Dβ1 RSS. All of these mice develop a greater percentage of αβ T cells expressing V2^+^ or V31^+^ TCRβ protein at the expense of cells using another type of TCRβ chain. We demonstrate that each Vβ RSS replacement increases Vβ rearrangement before feedback inhibition, competes with the homologous allele for usage in the TCRβ repertoire, and elevates the percentage of αβ T cells expressing TCRβ proteins from both alleles. We conclude that the poor qualities of Vβ RSSs for recombining with Dβ and Jβ RSSs enforce TCRβ allelic exclusion by stochastically limiting Vβ rearrangements before feedback inhibition from one allele halts further Vβ recombination.

## RESULTS

### Generation of Vβ RSS Replacement Mice with Grossly Normal αβ T Cell Development

To determine contributions of *Tcrb* RSSs in allelic exclusion, we established C57BL/6 mice carrying germline replacements of the *V2* or *V31* RSS with the stronger 3’Dβ1 RSS, referred to as the *V2^R^* or *V31^R^* modifications (Figures 1A, 1B, and S1C). We created mice with each replacement on one allele (*V2*^*R*/+^, *V31*^*R*/+^), both alleles (*V2^R/R^*, *V31^R/R^*), or opposite alleles (*V2^R^/V31^R^*). The assembly and expression of functional *Tcrb* genes is essential for αβ T cell development (Bouvier et al., 1996; Mombaerts et al., 1992). In thymocytes, Dβ-to-Jβ rearrangement initiates in ckit^+^CD25^-^ DN1 cells and continues in ckit^+^CD25^+^ DN2 and ckit^-^CD25^+^ DN3 cells, while Vβ-to-DJβ recombination initiates in DN3 cells (Godfrey and Zlotnik, 1993). The expression of a functional *Tcrb* gene in DN3 cells is necessary and rate-limiting for differentiation of ckit^-^CD25^-^ DN4 cells and then DP thymocytes (Baldwin et al., 2005; Serwold et al., 2007; Shinkai et al., 1992; Yang-Iott et al., 2010). We observed normal numbers and frequencies of splenic αβ T cells and thymocytes at each developmental stage in every genotype of Vβ RSS replacement mice (Figures 1C and S2A-S2F). The *V2^R^* and *V31^R^* alleles initiate *V2* and *V31* rearrangements in DN3 cells and at notably greater levels than *WT* alleles (Figures S3A-S3C). These data reveal that replacement of a *V2* and/or *V31* RSS with the stronger 3’Dβ1 RSS increases the frequency that Vβ recombination initiates without altering normal development or numbers of naïve αβ T cells.

### The 3’Dβ1 RSS-replaced Vβ Segments Outcompete Unmodified Vβs for Usage in the TCR Repertoire

In wild-type C57BL/6 mice, the representation of individual Vβ segments within the αβ TCR repertoires of DP thymocytes, SP thymocytes, and naïve splenic αβ T cells is similar and mirrors their relative levels of rearrangement in DN3 thymocytes (Wilson et al., 2001). Thus, we performed flow cytometry on mature naive αβ T cells (SP thymocytes and splenic αβ T cells) to determine effects of Vβ RSS substitutions on Vβ recombination and resultant usage in the αβ TCR repertoire. We used an antibody for a Cβ epitope contained in all TCRβ proteins in combination with different Vβ-specific antibodies that bind peptides encoded by a single Vβ [*V2, Trbv4 (V4), Trbv19 (V19)*, or *V31*] or a family of Vβs [*Trbv12.1* and *Trbv12.2* (*V12*) or *Trbv13.1, Trbv13.2*, and *Trbv13.3* (*V13*)]. In *WT* mice, we observed that 7.0% of SP cells express V2^+^ or V31^+^ TCRβ chains on their surface (Figures 1D-1F). For mice with *V2* or *V31* RSS replacement on one or both alleles, we detected a 6-11-fold increased representation of each modified Vβ on SP cells (Figures 1D-1F). Specifically, we detected V2^+^ TCRβ chains on 40.9% of cells from *V2*^*R*/+^ mice and on 61.4% of cells from *V2^R/R^* mice, and V31^+^ TCRβ chains on 50.0% of cells from *V31*^*R*/+^ mice and on 77.1% of cells from *V31^R/R^* mice (Figures 1D-1F). As all six genotypes exhibit similar numbers of SP cells (Figure 1C), the increased utilization of each RSS-replaced Vβ must be at the expense of other Vβ segments. Indeed, the percentages of V31^+^ SP cells are reduced in *V2*^*R*/+^ mice compared to *WT* mice (5.1% versus 7.0%) and in *V2^R/R^* mice relative to *V2*^*R*/+^ mice (3.8% versus 5.1%, Figures 1D-1F). Likewise, the percentages of V2^+^ SP cells are reduced in *V31*^*R*/+^ mice compared to *WT* mice (4.3% versus 7.0%) and in *V31^R/R^* mice relative to *V31*^*R*/+^ mice (2.3% versus 4.3%, Figures 1D-1F). Moreover, the percentage of SP cells expressing V4^+^, V12^+^, V13^+^, or V19^+^ TCRβ protein is lower than normal in *V2*^*R*/+^ and *V31*^*R*/+^ mice, and further reduced in *V2^R/R^* and *V31^R/R^* mice (Figures 1D, 1G, and data not shown). The Vβ usage in splenic αβ T cells of each Vβ RSS replacement mouse genotype is altered similarly as on SP thymocytes (Figures S4A-S4E). These data show that the stronger 3’Dβ1 RSS empowers *V2* and *V31* to outcompete normal Vβ segments for recombination and resultant usage in the αβ TCR repertoire.

Notably, each genotype of homozygous Vβ RSS replacement mice has a ~1.5-fold greater representation of its modified Vβ compared to the corresponding heterozygous genotype (Figure S4F). This less than additive effect based on allelic copy number suggests that *Tcrb* alleles compete for rearrangement and resultant usage in the αβ TCR repertoire. Our analysis of *V2*^*R*/+^, *V31*^*R*/+^, and *V2^R^/V31^R^* mice yields additional evidence for this competition as each RSS-replaced Vβ is less represented in *V2^R^*/*V31^R^* mice relative to *V2*^*R*/+^ or *V31*^*R*/+^ mice (Figures 1D-1F and S4A-S4C). Specifically, V2 is expressed on 32.2% of SP cells in *V2^R^/V31^R^* mice compared to 40.9% in *V2*^*R*/+^ mice, and V31 is expressed on 42.0% of SP cells in *V2^R^/V31^R^* mice relative to 50.0% in *V31*^*R*/+^ mice (Figures 1D-1F). We observed similar differences among splenic αβ T cells (Figures S4A-S4C). These differences imply that the overall Vβ recombination efficiency of each RSS-replaced allele is elevated such that it effectively competes with the other allele for recombination in thymocytes and usage in the αβ TCR repertoire.

Competition between *Tcrb* alleles implies that rearrangement of the unmodified allele in heterozygous Vβ RSS replacement mice might limit the extent that each RSS-replaced Vβ outcompetes other Vβ segments on the modified allele. To test this, we generated mice with the *WT*, *V2^R^*, or *V31^R^* allele opposite an allele with deletion of the *Tcrb* enhancer (Eβ). As Eβ deletion blocks all *Tcrb* recombination events *in cis* (Bories et al., 1996; Bouvier et al., 1996), an Eβ-deleted (*Eβ^Δ^*) allele cannot compete with an active *Tcrb* allele. We compared the Vβ repertoires of mature αβ T cells from *WT/Eβ^Δ^*, *V2^R^/Eβ^Δ^*, and *V31^R^/Eβ^Δ^* mice to cells from *WT*, *V2*^*R*/+^, and *V31*^*R*/+^ mice. The percentages of V2^+^ and V31^+^ SP thymocytes each are equivalent between *WT/Eβ^Δ^* and *WT* mice (Figures 2A-2D). In contrast, representation of each RSS-replaced Vβ is ~1.5-fold greater in *V2^R^*/*Eβ^Δ^* or *V31^R^/Eβ^Δ^* mice relative to *V2*^*R*/+^ or *V31*^*R*/+^ mice, respectively (Figures 2A-2D). Furthermore, the percentages of V2^+^ and V31^+^ cells in *V2^R^/Eβ^Δ^* and *V31^R^/Eβ^Δ^* mice are similar to those of *V2^R/R^* and *V31^R/R^* mice, respectively (compare Figures 1D-1F with Figures 2A-2D). These comparisons reveal that recombination of a wild-type *Tcrb* allele indeed limits the extent to which each RSS-replaced Vβ can outcompete other Vβ segments on the same allele.

**Figure 2.**
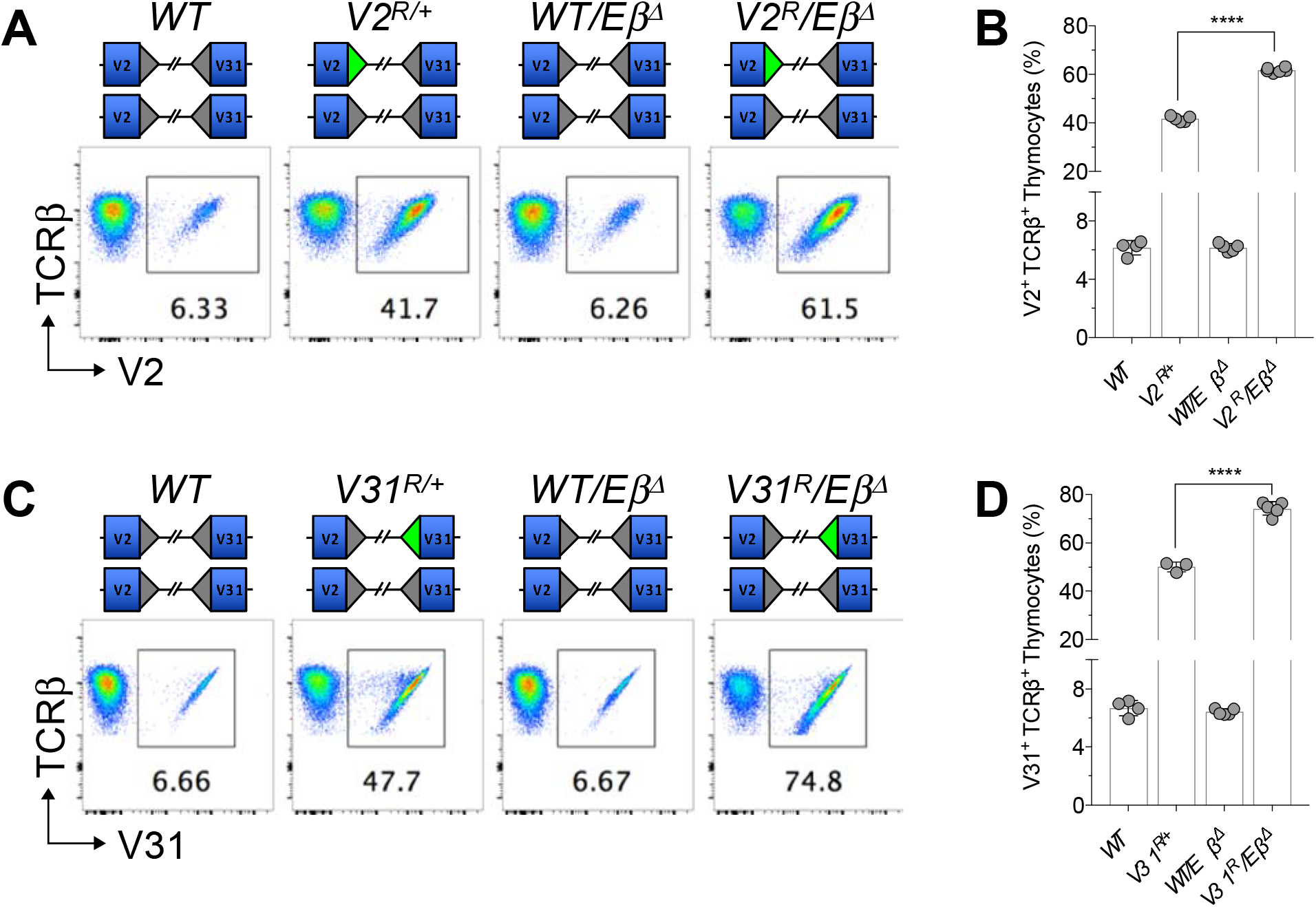
Vβ RSS Replacement Alleles Compete with Normal *Tcrb* Alleles for Usage in the TCRβ Repertoire. (A and C) Representative plots of SP thymocytes expressing V2^+^ (A) or V31^+^ (C) TCRβ chains. (B and D) Quantification of V2^+^ (B) or V31^+^ (D) SP thymocytes (mean ± SD, n ≥ 3, \*\*\*\**p*<0.0001).

### Vβ RSS Replacements Increase Biallelic Assembly and Expression of Functional TCRβ Genes

We next determined effects of Vβ RSS replacements on monoallelic TCRβ expression. Due to the absence of allotypic markers that can identify TCRβ chains from each allele, the field assays TCRβ allelic exclusion by quantifying cells that stain with two different anti-Vβ antibodies. This method suggests that 1-3% of αβ T cells exhibits biallelic TCRβ expression (Balomenos et al., 1995; Steinel et al., 2014). However, this might be an underestimation as antibodies are not available for all Vβ proteins and biallelic *Tcrb* expression involving the same Vβ segment cannot be discerned. Regardless, we used this approach to determine the percentages of αβ T cells expressing two different types of TCRβ chains in *WT*, *V2*^*R*/+^, *V2^R/R^*, *V31*^*R*/+^, *V31^R/R^*, and *V2^R^/V31^R^* mice. We first used an antibody for V2 or V31 combined with an antibody for V4, V12, V13, or V19. For each combination, we observed that 0.05-0.21% of SP cells stained with both antibodies in *WT* mice (Figures 3A-3D). In *V31*^*R*/+^ and *V31^R/R^* mice, we detected increased frequencies of SP cells that stained for V31 and each other Vβ tested (Figures 3C and 3D). Likewise, for *V2*^*R*/+^ and *V2^R/R^* mice, we saw increased frequencies of SP cells that stained for V2 and each other Vβ (Figures 3A and 3B). We also observed inverse trends where the frequencies of SP cells expressing V2 and another Vβ decreased in *V31*^*R*/+^ and *V31^R/R^* mice, as well as the frequencies of SP cells expressing V31 and another Vβ decreased in *V2*^*R*/+^ and *V2^R/R^* mice (Figures 3A-3D). We next quantified V2^+^V31^+^ cells and observed that 0.09% of SP thymocytes stained with both V2 and V31 antibodies in *WT* mice (Figures 3E and 3F). In mice carrying *V2^R^* or *V31^R^* on one or both alleles, we detected 0.3-0.68% of SP cells stained with both antibodies (Figures 3E and 3F). Strikingly, the frequency of V2^+^V31^+^ SP cells is 27-fold higher in *V2^R^/V31^R^* mice compared to *WT* mice (2.47% versus 0.09%, Figures 3E and 3F). To address any potential background staining from the increased frequencies of V2^+^ and V31^+^ cells in *V2^R^/V31^R^* mice, we mixed equal numbers of SP cells from *V2^R/R^* and *V31^R/R^* mice. Notably, the frequency of V2^+^V31^+^ cells in *V2^R^/V31^R^* mice is 3.5-fold greater than in mixed *V2^R/R^* and *V31^R/R^* cells (Figures 3E and 3F). This provides firm evidence that V2^+^V31^+^ cells in *V2^R^/V31^R^* mice are αβ T cells expressing both V2^+^ and V31^+^ TCRβ chains. The sum of the frequencies of double-staining cells for all Vβ combinations tested shows that the total incidence of SP cells expressing two types of TCRβ chains is increased for each Vβ RSS replacement genotype (Figure 3G). The highest incidence of dual-TCRβ expressing thymocytes is in *V2^R^/V31^R^* mice and is 5-fold more than in *WT* mice (Figure 3G). Similar increased incidences of dual-TCRβ expression was observed in splenic αβ T cells (Figures S5A-S5G). Collectively, these data provide strong evidence that replacement of a *V2* and/or *V31* RSS with the 3’Dβ1 RSS elevates the frequencies of mature αβ T cells exhibiting biallelic TCRβ expression.

**Figure 3.**
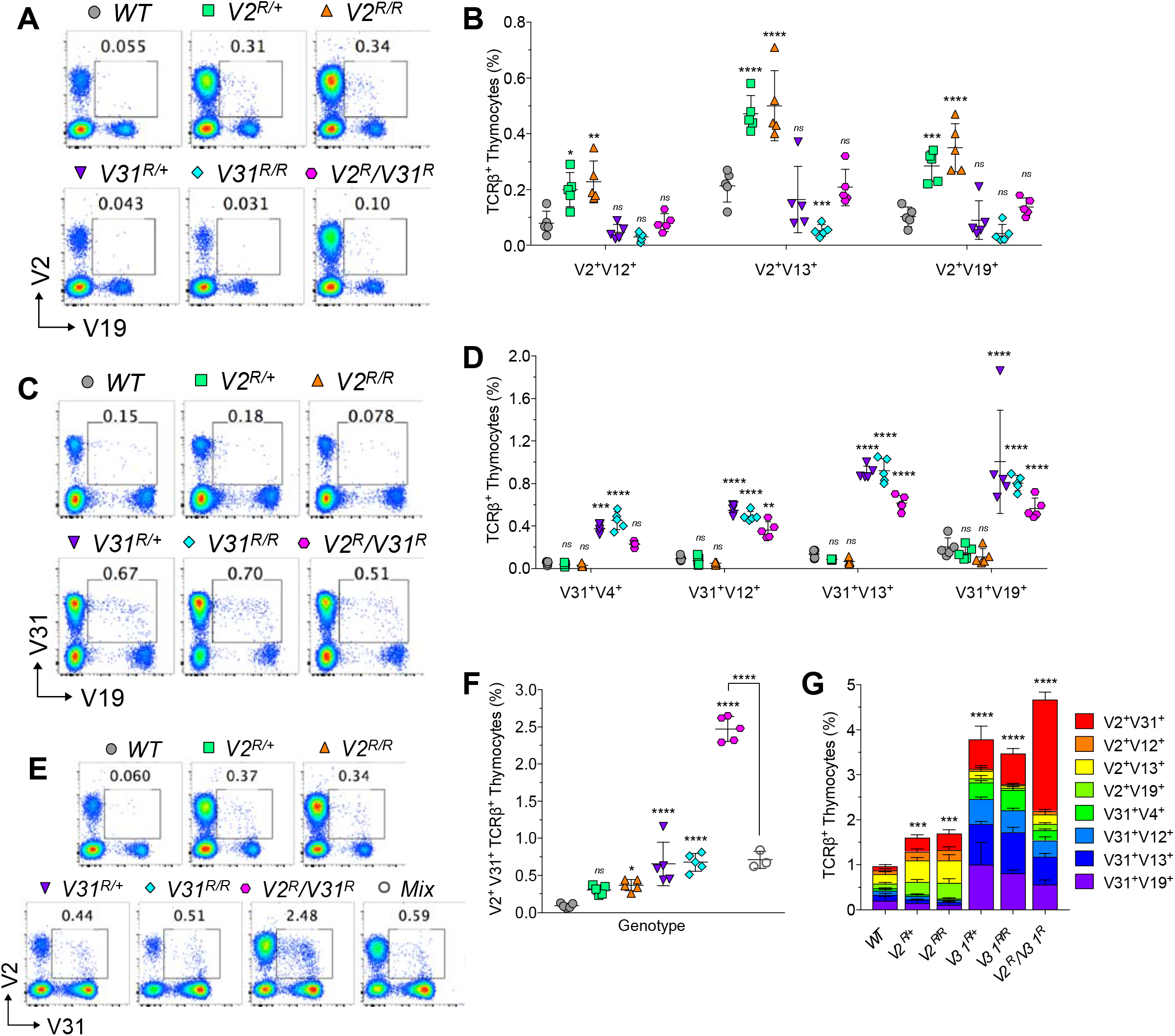
3’Dβ1 RSS-replaced Vβ Segments Increase Bi-allelic *Tcrb* Gene Expression. (A, C, and E) Representative plots of SP thymocytes expressing both V2^+^ and V19^+^ (A), V31^+^ and V19^+^ (B), or V2^+^ and V31^+^ (E) TCRβ chains. (B, D, and F) Quantification of SP thymocytes expressing the two indicated TCRβ chains. (F) *V2^R/R^* and *V31^R/R^* thymocytes were mixed 1:1 and analyzed. (B and D) n = 5, two-way ANOVA. (F) n ≥ 3, one-way ANOVA. (G) Quantification of double-staining SP thymocytes for each Vβ combination tested (n = 5, two-way ANOVA). All quantification plots show mean ± SD. Multiple post-tests are compared to *WT* unless indicated by bars, and *p*-values are corrected for multiple tests. ns=not significant, \**p*<0.05, \*\**p*<0.01, \*\*\**p*<0.001, \*\*\*\**p*<0.0001.

The elevated biallelic TCRβ expression must result from the 3’Dβ1 RSS increasing overall Vβ recombination so that both alleles assemble functional genes in a higher than normal percentage of thymocytes. To validate this, we made 102 αβ T cell hybridomas from splenocytes of *V2^R^/V31^R^* mice and analyzed *Tcrb* rearrangements by Southern blotting and PCR/sequencing. We compared our data to a prior study of 212 wild-type αβ T cell hybridomas, where 56.6% contained a single Vβ rearrangement on one allele and a DβJβ rearrangement(s) on the other allele, and 43.4% contained one in-frame and one out-of-frame Vβ rearrangement on opposite alleles (Table 1) (Khor and Sleckman, 2005). Of our *V2^R^/V31^R^* hybridomas, 45.1% had a single Vβ rearrangement on one allele and a DβJβ rearrangement on the other allele, while 31.4% carried one Vβ rearrangement on each allele (Table 1). We discovered that an additional 9.8% of *V2^R^/V31^R^* hybridomas had two Vβ rearrangements (involving *V31* and another Vβ) on one allele and a DβJβ rearrangement on the other allele, and another 13.7% harbored recombination of *V31* and another Vβ on one allele and a single Vβ rearrangement on the other allele (Table 1). Our data reveal a higher incidence of these rearrangements in *V2^R^/V31^R^* cells (23.5% versus 0%, Table 1) as none of the *WT* hybridomas had two distinct Vβ rearrangements on the same allele (Khor and Sleckman, 2005). In total, we observed monoallelic Vβ rearrangements in 54.9% of *V2^R^/V31^R^* hybridomas and biallelic Vβ rearrangements in 45.1% (Table 1). However, our sample size does not allow us to conclude that biallelic Vβ recombination is increased relative to the theoretical 40% or published 43.4% in *WT* αβ T cells (Figure S1A, Table 1) (Khor and Sleckman, 2005). Of the *V2^R^/V31^R^* hybridomas, 12.7% had a *V2* rearrangement and 56.9% had a *V31* rearrangement (Table 1), reflecting the increased usage of V2 and V31 in the TCR repertoire of *V2^R^/V31^R^* mice. Eight hybridomas (7.8% of total) had recombination of each RSS-replaced Vβ (Table 1). Although the 3’Dβ1 RSS is strongest at recombining to 5’Dβ RSSs, it also drives lower levels of recombination to Jβ RSSs (Banerjee and Schatz, 2014; Drejer-Teel et al., 2007; Jung et al., 2003; Tillman et al., 2003; Wu et al., 2003). *V31* rearrangement to a 5’Dβ RSS or Jβ RSS occurs by inversion and leaves a signal join in the locus, allowing for definitive identification of *V31* recombination to a DβJβ complex versus a Jβ segment by Southern blotting (Malissen et al., 1986; Wu et al., 2003). On *WT* alleles, *V31*-to-Jβ rearrangements have not been identified in 512 hybridomas spanning three studies (Bassing et al., 2000; Khor and Sleckman, 2005; Sleckman et al., 2000). Of the 58 *V31* rearrangements in *V2^R^/V31^R^* hybridomas, 33 were to a DβJβ complex, 24 were to a Jβ segment, and the target of one was undetermined (Tables 1 and 2). We sequenced the *V2* and *V31* rearrangements in the eight *V2^R^/V31^R^* hybridomas that recombined both *V2* and *V31*. Every *V2* rearrangement contained a Jβ2 segment and had one or more potential Dβ nucleotides (Table 2). For *V31*, we identified rearrangements to DβJβ complexes, the Vβ coding region of a *Trbv29-* Dβ1Jβ2 rearrangement, and the 5’Dβ1 RSS in a manner that produced a hybrid join (Table 2). Two hybridomas contained an in-frame *V2*DβJβ rearrangement on one allele and an in-frame *V31*DβJβ rearrangement on the other allele (Tables 1 and 2). Notably, this incidence of an in-frame Vβ(Dβ)Jβ rearrangement on each allele in ~2% of hybridomas mirrors the 2.47% of V2^+^V31^+^ αβ T cells in *V2^R^/V31^R^* mice. Our hybridoma data show that replacement of a single Vβ RSS with the 3’Dβ1 RSS on opposite alleles increases the frequency of overall Vβ recombination and biallelic assembly of functional *Tcrb* genes.

**Table 1.**
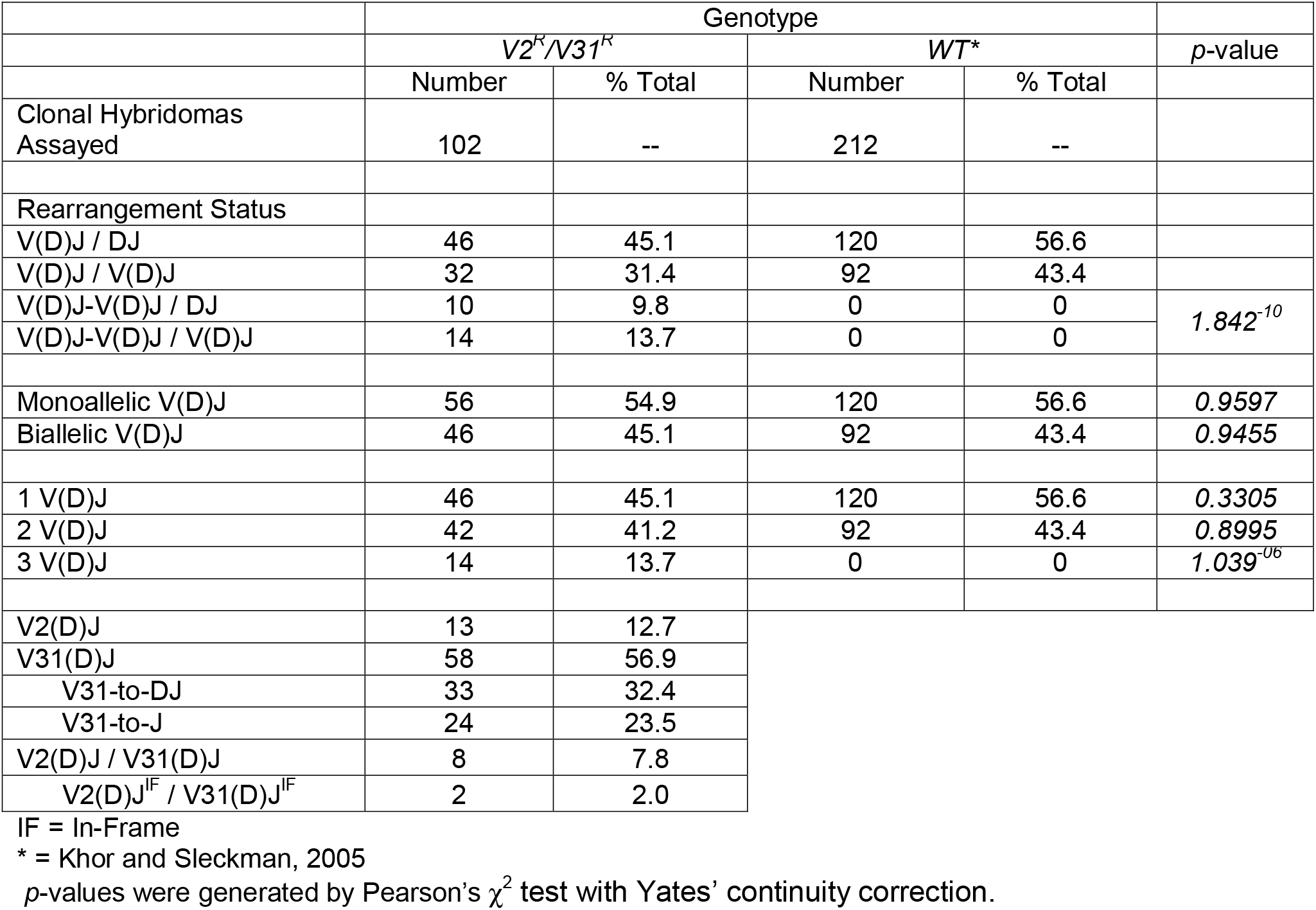
Analysis of Tcrb rearrangements in αβ T cell hybridomas

**Table 2.**
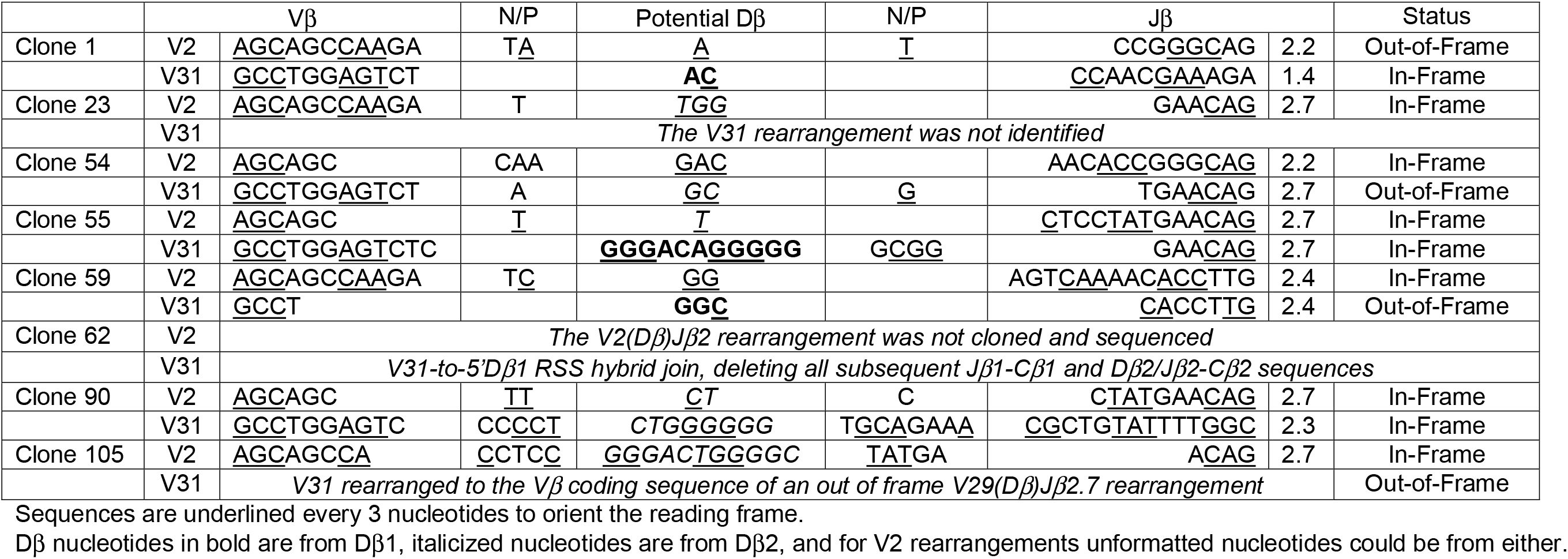
Sequence analysis of the *V2* and *V31* rearrangements on opposite alleles in *V2^R^/V31^R^* T cell hybridomas

### Vβ RSS Substitutions Elevate Vβ Recombination Independent of c-Fos Transcription Factor Binding

The increased Vβ recombination and TCRβ allelic inclusion in Vβ RSS replacement mice can be explained by the greater strength of the 3’Dβ1 RSS in recombining with Dβ and Jβ RSSs. Yet, this interpretation is complicated by the fact that 3’Dβ RSSs, but not Vβ RSSs, contain a c-Fos transcription factor binding site that spans the heptamer and spacer. *In vitro*, c-Fos interacts with RAG proteins when bound to 3’Dβ RSSs (Figure 4A) (Wang et al., 2008), leading to a model where c-Fos deposits RAG on 3’Dβ RSSs to sterically hinder Vβ recombination until Dβ-to-Jβ recombination deletes a 3’Dβ RSS (Wang et al., 2008). To investigate potential effects of c-Fos-mediated RAG deposition and/or transcription-associated accessibility, we made C57BL/6 mice carrying germline *V2* or *V31* RSS replacements with a 3’Dβ1 variant RSS, referred to as the *V2^F^* or *V31^F^* modification. The 3’Dβ1 RSS variant contains a two-nucleotide substitution in the spacer that abolishes both c-Fos binding and c-Fos-mediated RAG deposition, but has no obvious effect on the activity of the 3’Dβ1 RSS at recombining to a V_κ_ RSS *in vitro* (Figures 4A and 4B) (Wang et al., 2008). We established mice with each Vβ RSS replacement on one allele (*V2*^*F*/+^ and *V31*^*F*/+^) or opposite alleles (*V2^F^/V31^F^*).

**Figure 4.**
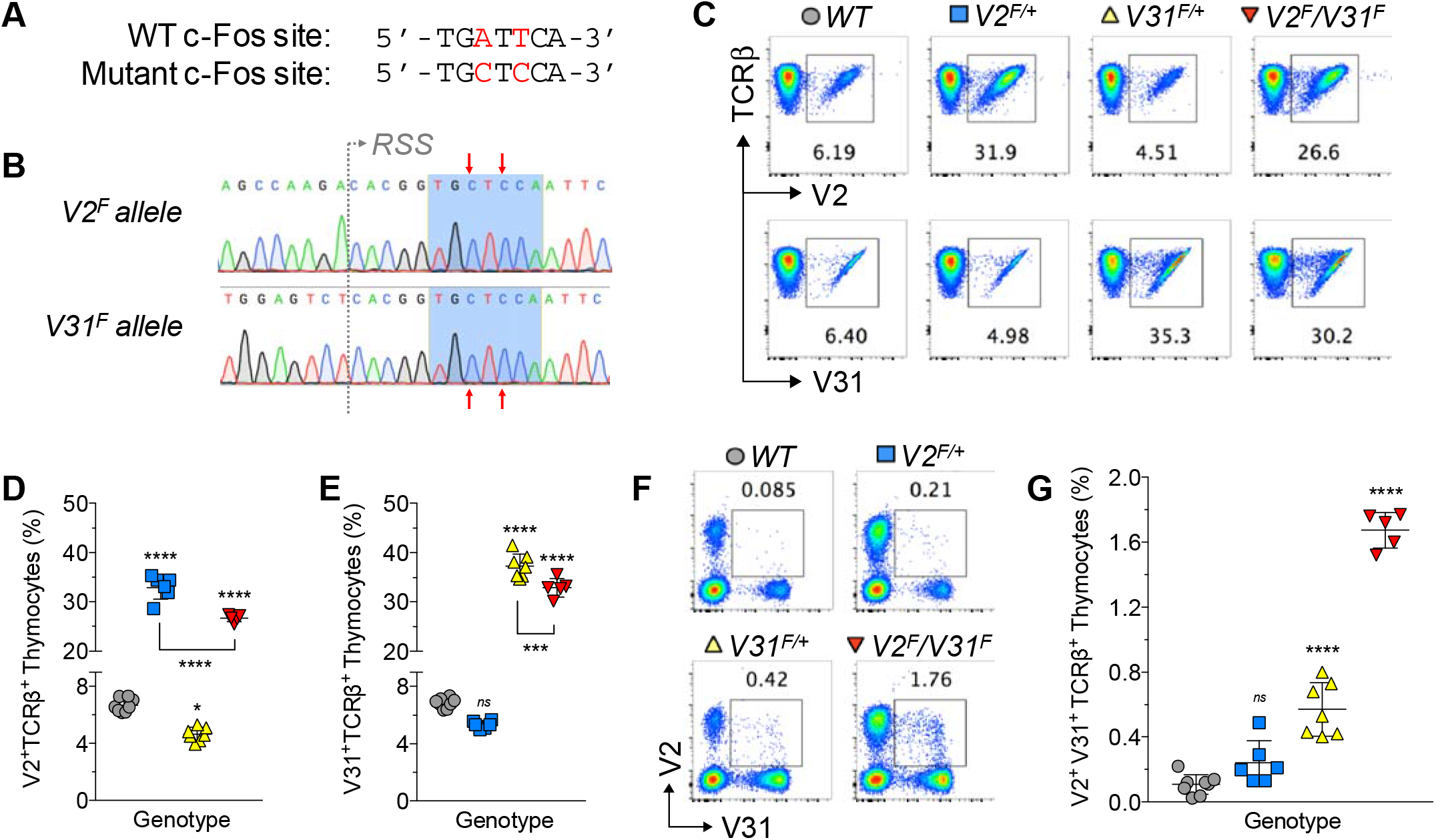
3’Dβ1 RSS Substitutions Increase Vβ usage and TCRβ Allelic Inclusion Independent of c-Fos binding. (A) Sequences of the normal and inactivated c-Fos binding site in the 3’Dβ1 RSS. The A→C and T→C mutations are indicated in red. (B) Sequence validation of the V2 or V31 RSS replacement with the variant 3’Dβ1 RSS. The inactivated c-Fos binding site is highlighted in blue and mutated nucleotides indicated by red arrows. (C) Representative plots of SP thymocytes expressing V2^+^ or V31^+^ TCRβ chains. (D and E) Quantification of V2^+^ (D) or V31^+^ (E) SP thymocytes (n ≥ 5, one-way ANOVA). (F) Representative plots of SP thymocytes expressing both V2^+^ and V31^+^ TCRβ chains. (G) Quantification of SP thymocytes expressing both V2^+^ and V31^+^ TCRβ chains (n ≥ 5, one-way ANOVA). All quantification plots show mean ± SD. Multiple post-tests are compared to *WT* and *p*<-values are corrected for multiple tests. ns=not significant, \**p*<0.05, \*\*\*\**p*<0.0001.

We performed flow cytometry on *WT*, *V2*^*F*/+^, *V31*^*F*/+^, and *V2^F^/V31^F^* mice to determine effects of *V2^F^* and/or *V31^F^* alleles on αβ T cell development, TCRβ repertoire, and TCRβ allelic exclusion. The numbers and frequencies of splenic αβ T cells and thymocytes at each developmental stage are normal in each Vβ RSS replacement genotype (data not shown), indicating no major effects on αβ T cell development. In contrast, replacement of the *V2* and/or *V31* RSS with the 3’Dβ1 RSS variant increases the frequencies of SP cells expressing V2, V31, or both (Figures 4C-4E). Specifically, we found a 4.9-fold greater than normal representation of V2^+^ cells in *V2^F/+^* mice and a 5.4-fold higher than normal frequency of V31^+^ cells in *V31^F/+^* mice (Figures 4C-4E). Similar to what we observed in *V2^R^/V31^R^* mice, the *V2^F^* and *V31^F^* alleles compete for recombination in thymocytes as we detected smaller increases of V2^+^ and V31^+^ cells in *V2^F^/V31^F^* mice (Figures 4C-4E). Finally, we detected higher than normal frequencies of V2^+^V31^+^ SP cells in *V2*^*F*/+^, *V31*^*F*/+^, and *V2^F^/V31^F^* mice (Figures 4F and 4G), revealing that replacement of a *V2* and/or *V31* RSS with the 3’Dβ1 RSS variant elevates the incidence of biallelic TCRβ expression. These data indicate that neither RAG deposition nor transcription-associated accessibility from potential c-Fos binding is needed for 3’Dβ1 RSS substitutions to increase Vβ rearrangement and biallelic TCRβ expression.

### Vβ RSS Replacements Increase the Initiation of Vβ Recombination before Feedback Inhibition

The elevated incidences of biallelic TCRβ expression in Vβ RSS replacement mice could arise from increased initiation of Vβ recombination prior to enforcement of feedback inhibition and/or continued Vβ recombination during feedback inhibition. We found that the rearrangements of RSS-replaced Vβs are elevated in DN3 thymocytes of *V2^R^/Eβ^Δ^* and *V31^R^/Eβ^Δ^* mice (Figure S3C). As these Vβs rearrange independent of competition and feedback inhibition from the other allele, the 3’Dβ1 RSS substitution does increase Vβ recombination before feedback inhibition. To determine if rearrangements of RSS-replaced Vβs are halted by TCRβ-mediated feedback inhibition, we established and analyzed *V2*^*R*/+^ and *V31*^*R*/+^ mice with a pre-assembled functional TCRβ transgene (*Tcrb^Tg^*). Expression of the transgene-encoded V13^+^ TCRβ chain initiates in DN2 cells and signals feedback inhibition of Vβ rearrangements (Steinel et al., 2010). However, ~3% of *Tcrb^Tg^* αβ T cells also expresses TCRβ protein from VβDβJβ rearrangements that occur in DN3 cells before *Tcrb^Tg^*-mediated feedback inhibition (Steinel et al., 2010). We observed that the *Tcrb^Tg^* more effectively decreases utilization of *V2* than *V31* when each is flanked by their own RSS or the 3’Dβ1 RSS (Figures 5A-5D). Next, we made αβ T cell hybridomas from *V2^R^*^/+^, *Tcrb^Tg^V2*^*R*/+^, *V31*^*R*/+^, and *Tcrb^Tg^V31*^*R*/+^ mice to quantify Vβ rearrangements. We detected *V2* rearrangements in 50% of *V2*^*R*/+^ clones but not in any *Tcrb^Tg^V2*^*R*/+^ cells (*p* = 2.68×10^-5^, Pearson’s χ^2^ test with Yates’ correction), and we found *V31* rearrangements in 50% of *V31*^*R*/+^ clones and in only 15% of *Tcrb^Tg^V31*^*R*/+^ cells (*p* = 1.63×10^-5^, Pearson’s χ^2^ test with Yates’ correction, Table S1). Our previous analysis of 129 *Tcrb^Tg^* αβ T cell hybridomas showed that 2.3% had a *V31* rearrangement and an additional 7% carried recombination of a different Vβ (Table S1 and data not shown) (Steinel et al., 2010). Collectively, these data indicate that *Tcrb^Tg^*-signaled feedback inhibition suppresses recombination of RSS-replaced Vβs and does so more effectively for *V2^R^* versus *V31^R^*. As feedback inhibition could block accessibility of 5’Dβ RSSs in DN cells (Bassing et al., 2000), RSS-replaced Vβs may continue to recombine with Jβ segments until Vβ recombination is silenced by differentiation of DP thymocytes. To assess this possibility, we quantified *V31* targeting to DβJβ complexes or Jβ segments in *V31*^*R*/+^ and *Tcrb^Tg^V31*^*R*/+^ hybridomas where *V31* is the only rearranged Vβ segment. These primary *V31* rearrangements occurred to Jβ segments in 38% of *V31*^*R*/+^ and 14% of *Tcrb^Tg^V31*^*R*/+^ cells (Table S1), revealing that TCRβ-mediated feedback inhibition also suppresses *V31^R^*-to-Jβ recombination. Our hybridoma analysis reveals that increased initiation of Vβ recombination prior to TCRβ-mediated feedback inhibition is the predominant, if not sole, mechanistic basis for the elevated frequencies of biallelic TCRβ expression in Vβ RSS replacement mice.

**Figure 5.**
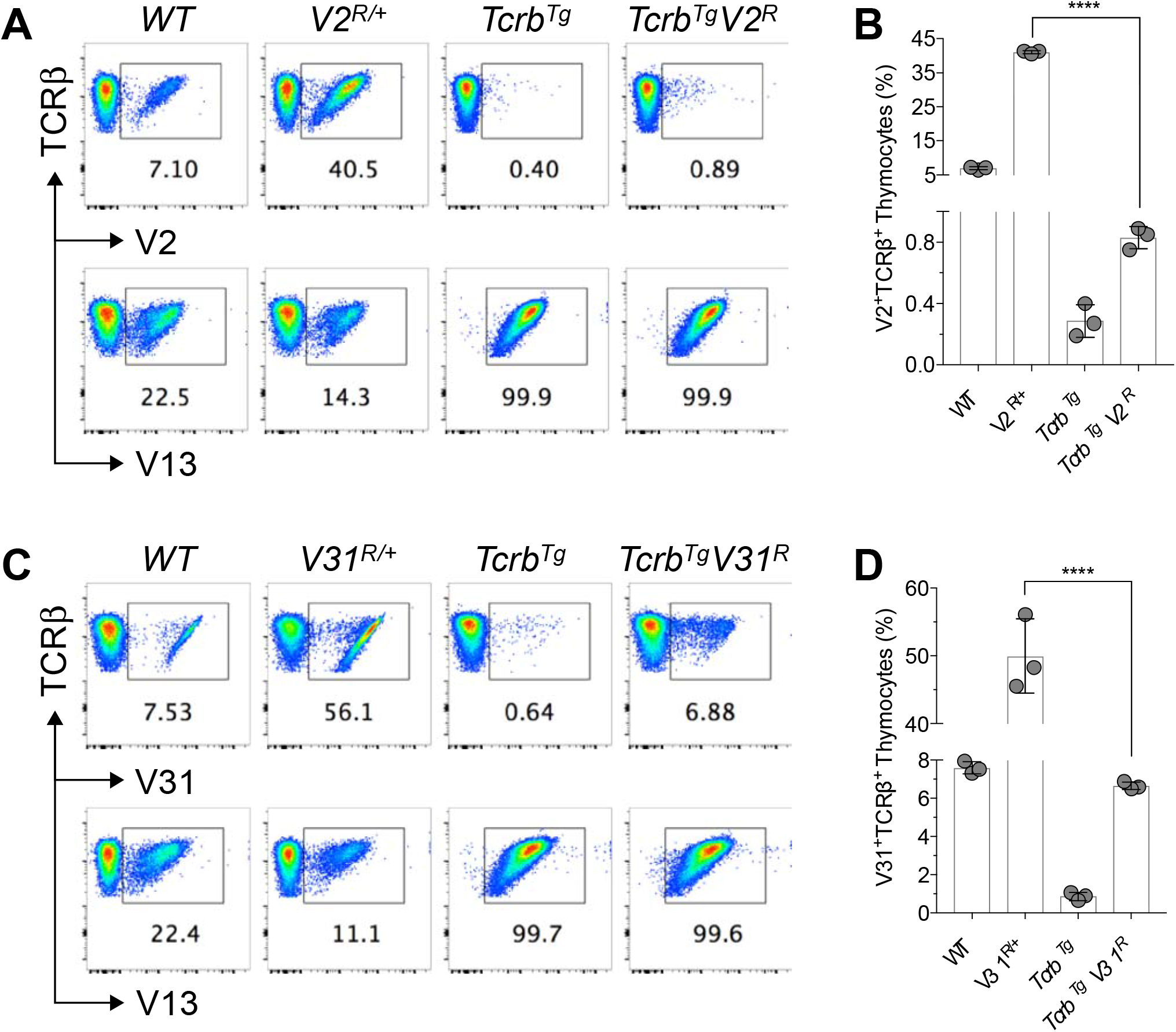
3’Dβ1 RSS Substitutions Increase Vβ recombination before Enforcement of Feedback Inhibition. (A and C) Representative plots of SP thymocytes expressing V2^+^ (A), V31^+^ (C), or V13^+^ TCRβ chains. (B and D) Quantification of SP thymocytes expressing V2^+^ (B) or V31^+^ (D) TCRβ chains (n = 3, \*\*\*\**p*<0.0001). All quantification plots show mean ± SD.

## DISCUSSION

### Improving the Quality of Vβ RSSs Increases Vβ Rearrangement and Biallelic TCRβ Expression

Here we show that elevating Vβ rearrangement frequency by replacing the endogenous *V2* and/or *V31* RSS with the 3’Dβ1 RSS increases the incidences of biallelic TCRβ expression before TCRβ-signaled feedback inhibition. Nucleotide sequence differences between the 3’Dβ1 and *V2* or *V31* RSSs must provide the mechanistic basis for enhanced Vβ rearrangement. Although the 3’Dβ1 RSS can bind c-Fos to recruit RAG *in vitro* (Wang et al., 2008), we demonstrate that a variant 3’Dβ1 RSS that cannot bind c-Fos maintains elevated levels of Vβ recombination and biallelic TCRβ expression *in vivo*. The increased rearrangement and preferential targeting of RSS-replaced *V2* and *V31* to DJβ complexes mirrors the relative *in vitro* activities of the 3’Dβ1, *V2*, and *V31* RSSs for RAG/HMGB1-mediated synapsis and cleavage (Banerjee and Schatz, 2014; Drejer-Teel et al., 2007; Jung et al., 2003). Context dependent kinking of each RSS spacer must occur for RAG/HMGB1 to pair RSSs for coupled cleavage (Kim et al., 2018). Indeed, the normal and variant 3’Dβ1 RSSs possess spacers of greater bending quality compared to the *V2* and *V31* RSSs (Kim et al., 2018) and heptamers and nonamers of less overall variation from consensus (Figure 1B). Accordingly, the increased levels of Vβ rearrangement in our RSS replacement mice can be explained mechanistically by the greater quality of the 3’Dβ1 RSS for RAG/HMGB1-mediated pairing and coupled cleavage with 5’Dβ and Jβ RSSs. Although our observations cannot exclude potentially minor contributions of RSS differences in regulating RAG cleavage in Vβ chromatin, our 3’Dβ1 RSS replacements neither introduce a recognized transcription factor binding site nor increase germline transcription of *V2* or *V31* (data not shown). Some RSSs, albeit not the 3’Dβ1 RSS, position nucleosomes over themselves even within accessible chromatin (Baumann et al., 2003; Kondilis-Mangum et al., 2010). Thus it is possible that Vβ RSSs, but not 3’Dβ1 RSSs, bind nucleosomes to antagonize recombination of otherwise accessible Vβ segments since RSS nucleosome occupancy inhibits RAG access and cleavage (Golding et al., 1999; Kwon et al., 1998). Even so, improving the quality of a Jβ RSS in mice lacking Dβ segments permits robust Vβ-to-Jβ rearrangements where the normal poor quality Vβ RSSs are intact (Bassing et al., 2000). Therefore, we conclude that the improved Vβ RSS quality for recombining with Dβ and Jβ RSSs is the underlying mechanism for increased Vβ recombination and biallelic TCRβ expression in our Vβ RSS replacement mice.

### The Poor Qualities of Vβ RSSs Provide a Stochastic Mechanism for Limiting Vβ Recombination Before Feedback Inhibition

Our study offers the first validated mechanism for how only one allele of any AgR locus assembles a functional gene before feedback inhibition. In addition to promoting Vβ recombination and biallelic TCRβ expression, improving the quality of one Vβ RSS reveals that *Tcrb* alleles compete for Vβ recombination. The only way that elevating the efficiency of Vβ recombination on one allele can outcompete Vβ rearrangement on the other is if both alleles are activated in the time window(s) that feedback inhibition requires to block Vβ recombination of the second allele. If deterministic models were correct, the recombination of one allele would have no consequence on the recombination of the other, and one would predict that the increases of Vβ recombination in RSS replacement mice would be additive based on allelic copy number. Consequently, our data demonstrate that the poor qualities of Vβ RSSs for recombining with Dβ and Jβ RSSs provides a stochastic mechanism that serves a major role in limiting the incidence of functional Vβ rearrangements on both alleles before feedback inhibition terminates Vβ recombination.

We propose the following model for how thymocytes enforce TCRβ allelic exclusion. In non-cycling DN3 cells, at least one allele becomes active and undergoes Vβ-to-DJβ recombination. The DNA DSBs trigger transient feedback inhibition at least in part by repression of RAG expression, providing time to test the initial rearrangement (Fisher et al., 2017; Steinel et al., 2014). In cells where both alleles are activated, poor Vβ RSSs limit the likelihood of Vβ recombination on both alleles before loss of RAG. If the rearrangement is out-of-frame, RAG re-expression permits Vβ recombination on the second allele, or on the first allele if a Dβ2Jβ2 complex is available. In the latter case, poor Vβ RSSs again decrease the chance for Vβ recombination on both alleles. When the first rearrangement is in-frame, its TCRβ protein activates Cyclin D3 to move cells into S phase (Sicinska et al., 2003), where RAG2 is degraded (Lin and Desiderio, 1994).

Based on its function in pro-B cells (Powers et al., 2012), Cyclin D3 may repress Vβ accessibility before cells enter S phase. In DN3 cells where RAG is re-expressed between DSB repair and S phase entry, poor Vβ RSSs limit the possibility of further Vβ recombination on the second allele. Additional factors, including inhibition of RC formation, Vβ accessibility, and locus contraction via stochastic interaction of alleles with nuclear lamina (Chan et al., 2013; Chen et al., 2018; Schlimgen et al., 2008), cooperate with poor Vβ RSSs to limit biallelic assembly as DN3 cells attempt *Tcrb* recombination. Finally, TCRβ signals promote genetic and epigenetic changes that silence Vβ recombination in DP cells where *Tcra* genes assemble (Jackson and Krangel, 2005; Liang et al., 2002; Majumder et al., 2015; Skok et al., 2007). Notably, all features of this model could apply to IgH allelic exclusion.

### The Broader Impacts of Vβ RSSs Being a Major Determinant of Vβ Recombination Frequency

The field has strived to elucidate mechanisms that promote V rearrangements across large chromosomal distances, with emphasis on factors that determine broad usage of V segments or promote allelic exclusion. *In vivo* experiments have shown that modulation of accessibility and RC contact of a V segment can influence its rearrangement frequency (Fuxa et al., 2004; Jain et al., 2018; Ryu et al., 2004). Correlative computational analyses conclude that V accessibility is the predominant factor for determining relative V utilization at *Tcrb* and *Igh* loci, while V RSS quality and RC contact each mainly function as a binary switch to prevent or allow recombination (Bolland et al., 2016; Gopalakrishnan et al., 2013). On the contrary, our data show that the qualities of RSSs flanking *V2* and *V31* function far beyond reaching a minimal threshold for functional synapsis and cleavage with Dβ RSSs. The increased usage of *V2^R^* and *V31^R^* at the expense of other Vβs on the same allele indicate that most, if not all, Vβs dynamically compete with each other for productive contact with the RC. On a normal allele, RAG bound to Dβ RSSs likely repeatedly capture and release different Vβ RSSs before synapsis proceeds to functional cleavage (Wu et al., 2003). This sampling of V segments could occur via diffusional-based synapsis of V RSSs positioned within a cloud of spatial proximity (Ji et al., 2010) or by RAG chromosomal loop scanning-based synapsis (Hu et al., 2015; Jain et al., 2018). To determine RSS quality (strength), the field typically uses an algorithm that calculates a recombination information content (RIC) score, which is based on statistical modeling of how each nucleotide diverges from an averaged RSS (Cowell et al., 2003). The RIC scores of the RSSs we manipulated predict that the 3’Dβ1 RSS replacement would decrease *V2* recombination and the variant 3’Dβ1 RSS substitution would reduce both *V2* and *V31* rearrangements (Figure S5H). These differences between predicted and empirical data could be due to a number of possibilities, including that the RIC algorithm does not address pairwise effects of RSSs on recombination. Regardless, the discrepancies between outcomes predicted by machine-generated associations and our actual *in vivo* results highlight the critical need to experimentally test computationally-based models of V(D)J recombination.

Since the discovery of AgR allelic exclusion (Pernis et al., 1965), the field has worked to identify mechanisms and physiological roles for monoallelic expression of TCR and Ig genes. The predominant, long-standing theory is that the expression of only one type (specificity) of AgR by T and B cells suppresses autoimmunity by ensuring negative selection of cells expressing a self-reactive receptor (Brady et al., 2010a; Heath and Miller, 1993; Padovan et al., 1993). Consistent with this hypothesis, expression of a second endogenous AgR enables cells bearing a transgenic autoreactive AgR to evade negative selection (Auger et al., 2012; Iliev et al., 1994; Sarukhan et al., 1998; Zal et al., 1996), and biallelic TCRα expression potentiates autoimmune diabetes in the NOD mouse model (Schuldt et al., 2017). Our findings suggest new avenues for investigation. As Vβ RSSs share sequence features with V_H_ RSSs, but not V RSSs of other loci (Liang et al., 2002), experimentally determining if V_H_ RSSs are poor for recombining with D_H_ RSSs to thus restrain V_H_ recombination and enforce IgH allelic exclusion is warranted. In addition, RSS replacements could test the model that poor activities of Igλ RSSs, relative to Igκ RSSs, help mediate isotypic exclusion (Ramsden and Wu, 1991) where most B cells express Igκ or Igλ, but not both types of light chains (Bernier and Cebra, 1964). *V2^R^/V31^R^* mice and the RSS replacement approach provide unprecedented experimental means to determine the effects of biallelic expression of diverse TCRβ chains in the αβ T cell population. RSS replacement mice also permit testing of possible additional reasons for controlling V rearrangements between alleles, including that monospecificity facilitates robust lymphocyte activation upon antigen encounter (Vettermann and Schlissel, 2010) and our view that asynchronous V-to-(D)J recombination between alleles suppresses RAG-triggered genomic instability and resultant lymphoid cancers.

## ACKNOWLEDGEMENTS

We thank Adele Harman and Jennifer Dunlap of the Children’s Hospital of Philadelphia (CHOP) Transgenics Core for help establishing all Vβ RSS replacement mice and the CHOP Flow Cytometry Core for assistance with analytic analyses and cell sorting. We thank Drs. Grace Teng, Yuhang Zhang, and David G. Schatz for helpful discussions of this research and the manuscript. The National Institutes of Health grants T32 AI055428 (G.S.W.) and RO1 AI 130231 (C.H.B.) supported this work.

## AUTHOR CONTRIBUTIONS

C.H.B. conceived and supervised this study. C.H.B. and K.S.Y.I. designed the *V2^R^* and *V31^R^* modifications. G.S.W. designed the *V2^F^* and *V31^F^*modifications. G.S.W. and C.H.B. designed the research plan. G.S.W., with assistance from K.D.L., conducted and analyzed all mouse experiments. K.S.Y.I. and M.A.R. made and analyzed hybridomas, and worked with G.S.W. and C.H.B. to identify *Tcrb* rearrangements. K.E.H. performed all statistical analyses. G.S.W. and C.H.B. worked together to prepare the manuscript.

## References

Akira, S., Okazaki, K., and Sakano, H. (1987). Two pairs of recombination signals are sufficient to cause immunoglobulin V-(D)-J joining. Science 238, 1134–1138.

Auger, J.L., Haasken, S., Steinert, E.M., and Binstadt, B.A. (2012). Incomplete TCR-beta allelic exclusion accelerates spontaneous autoimmune arthritis in K/BxN TCR transgenic mice. Eur J Immunol 42, 2354–2362.

Baldwin, T.A., Sandau, M.M., Jameson, S.C., and Hogquist, K.A. (2005). The timing of TCR alpha expression critically influences T cell development and selection. J Exp Med 202, 111–121.

Balomenos, D., Balderas, R.S., Mulvany, K.P., Kaye, J., Kono, D.H., and Theofilopoulos, A.N. (1995). Incomplete T cell receptor V beta allelic exclusion and dual V beta-expressing cells. J Immunol 155, 3308–3312.

Banerjee, J.K., and Schatz, D.G. (2014). Synapsis alters RAG-mediated nicking at Tcrb recombination signal sequences: implications for the “beyond 12/23” rule. Mol Cell Biol 34, 2566–2580.

Bassing, C.H., Alt, F.W., Hughes, M.M., D’Auteuil, M., Wehrly, T.D., Woodman, B.B., Gartner, F., White, J.M., Davidson, L., and Sleckman, B.P. (2000). Recombination signal sequences restrict chromosomal V(D)J recombination beyond the 12/23 rule. Nature 405, 583–586.

Bassing, C.H., Swat, W., and Alt, F.W. (2002). The mechanism and regulation of chromosomal V(D)J recombination. Cell 109 Suppl, S45–55.

Baumann, M., Mamais, A., McBlane, F., Xiao, H., and Boyes, J. (2003). Regulation of V(D)J recombination by nucleosome positioning at recombination signal sequences. EMBO J 22, 5197–5207.

Bernier, G.M., and Cebra, J.J. (1964). Polypeptide Chains of Human Gamma-Globulin: Cellular Localization by Fluorescent Antibody. Science 144, 1590–1591.

Bolland, D.J., Koohy, H., Wood, A.L., Matheson, L.S., Krueger, F., Stubbington, M.J., Baizan-Edge, A., Chovanec, P., Stubbs, B.A., Tabbada, K., et al. (2016). Two Mutually Exclusive Local Chromatin States Drive Efficient V(D)J Recombination. Cell Rep 15, 2475–2487.

Bories, J.C., Demengeot, J., Davidson, L., and Alt, F.W. (1996). Gene-targeted deletion and replacement mutations of the T-cell receptor beta-chain enhancer: the role of enhancer elements in controlling V(D)J recombination accessibility. Proc Natl Acad Sci U S A 93, 7871–7876.

Bouvier, G., Watrin, F., Naspetti, M., Verthuy, C., Naquet, P., and Ferrier, P. (1996). Deletion of the mouse T-cell receptor beta gene enhancer blocks alphabeta T-cell development. Proc Natl Acad Sci U S A 93, 7877–7881.

Brady, B.L., Oropallo, M.A., Yang-Iott, K.S., Serwold, T., Hochedlinger, K., Jaenisch, R., Weissman, I.L., and Bassing, C.H. (2010a). Position-dependent silencing of germline Vss segments on TCRss alleles containing preassembled VssDJssCss1 genes. J Immunol 185, 3564–3573.

Brady, B.L., Steinel, N.C., and Bassing, C.H. (2010b). Antigen receptor allelic exclusion: an update and reappraisal. J Immunol 185, 3801–3808.

Chan, E.A., Teng, G., Corbett, E., Choudhury, K.R., Bassing, C.H., Schatz, D.G., and Krangel, M.S. (2013). Peripheral subnuclear positioning suppresses Tcrb recombination and segregates Tcrb alleles from RAG2. Proc Natl Acad Sci U S A 110, E4628–4637.

Chen, S., Luperchio, T.R., Wong, X., Doan, E.B., Byrd, A.T., Roy Choudhury, K., Reddy, K.L., and Krangel, M.S. (2018). A Lamina-Associated Domain Border Governs Nuclear Lamina Interactions, Transcription, and Recombination of the Tcrb Locus. Cell Rep 25, 1729–1740 e1726.

Connor, A.M., Fanning, L.J., Celler, J.W., Hicks, L.K., Ramsden, D.A., and Wu, G.E. (1995). Mouse VH7183 recombination signal sequences mediate recombination more frequently than those of VHJ558. J Immunol 155, 5268–5272.

Cowell, L.G., Davila, M., Yang, K., Kepler, T.B., and Kelsoe, G. (2003). Prospective estimation of recombination signal efficiency and identification of functional cryptic signals in the genome by statistical modeling. J Exp Med 197, 207–220.

Drejer-Teel, A.H., Fugmann, S.D., and Schatz, D.G. (2007). The beyond 12/23 restriction is imposed at the nicking and pairing steps of DNA cleavage during V(D)J recombination. Mol Cell Biol 27, 6288–6299.

Farago, M., Rosenbluh, C., Tevlin, M., Fraenkel, S., Schlesinger, S., Masika, H., Gouzman, M., Teng, G., Schatz, D., Rais, Y., et al. (2012). Clonal allelic predetermination of immunoglobulin-kappa rearrangement. Nature 490, 561–565.

Fisher, M.R., Rivera-Reyes, A., Bloch, N.B., Schatz, D.G., and Bassing, C.H. (2017). Immature Lymphocytes Inhibit Rag1 and Rag2 Transcription and V(D)J Recombination in Response to DNA Double-Strand Breaks. J Immunol 198, 2943–2956.

Fuxa, M., Skok, J., Souabni, A., Salvagiotto, G., Roldan, E., and Busslinger, M. (2004). Pax5 induces V-to-DJ rearrangements and locus contraction of the immunoglobulin heavy-chain gene. Genes Dev 18, 411–422.

Gauss, G.H., and Lieber, M.R. (1992). The basis for the mechanistic bias for deletional over inversional V(D)J recombination. Genes Dev 6, 1553–1561.

Glusman, G., Rowen, L., Lee, I., Boysen, C., Roach, J.C., Smit, A.F., Wang, K., Koop, B.F., and Hood, L. (2001). Comparative genomics of the human and mouse T cell receptor loci. Immunity 15, 337–349.

Godfrey, D.I., and Zlotnik, A. (1993). Control points in early T-cell development. Immunol Today 14, 547–553.

Golding, A., Chandler, S., Ballestar, E., Wolffe, A.P., and Schlissel, M.S. (1999). Nucleosome structure completely inhibits in vitro cleavage by the V(D)J recombinase. EMBO J 18, 3712–3723.

Gopalakrishnan, S., Majumder, K., Predeus, A., Huang, Y., Koues, O.I., Verma-Gaur, J., Loguercio, S., Su, A.I., Feeney, A.J., Artyomov, M.N., et al. (2013). Unifying model for molecular determinants of the preselection Vbeta repertoire. Proc Natl Acad Sci U S A 110, E3206–3215.

Heath, W.R., and Miller, J.F. (1993). Expression of two alpha chains on the surface of T cells in T cell receptor transgenic mice. J Exp Med 178, 1807–1811.

Hesse, J.E., Lieber, M.R., Mizuuchi, K., and Gellert, M. (1989). V(D)J recombination: a functional definition of the joining signals. Genes Dev 3, 1053–1061.

Hewitt, S.L., Yin, B., Ji, Y., Chaumeil, J., Marszalek, K., Tenthorey, J., Salvagiotto, G., Steinel, N., Ramsey, L.B., Ghysdael, J., et al. (2009). RAG-1 and ATM coordinate monoallelic recombination and nuclear positioning of immunoglobulin loci. Nat Immunol 10, 655–664.

Horowitz, J.E., and Bassing, C.H. (2014). Noncore RAG1 regions promote Vbeta rearrangements and alphabeta T cell development by overcoming inherent inefficiency of Vbeta recombination signal sequences. J Immunol 192, 1609–1619.

Hu, J., Zhang, Y., Zhao, L., Frock, R.L., Du, Z., Meyers, R.M., Meng, F.L., Schatz, D.G., and Alt, F.W. (2015). Chromosomal Loop Domains Direct the Recombination of Antigen Receptor Genes. Cell 163, 947–959.

Iliev, A., Spatz, L., Ray, S., and Diamond, B. (1994). Lack of allelic exclusion permits autoreactive B cells to escape deletion. J Immunol 153, 3551–3556.

Jackson, A.M., and Krangel, M.S. (2005). Allele-specific regulation of TCR beta variable gene segment chromatin structure. J Immunol 175, 5186–5191.

Jain, S., Ba, Z., Zhang, Y., Dai, H.Q., and Alt, F.W. (2018). CTCF-Binding Elements Mediate Accessibility of RAG Substrates During Chromatin Scanning. Cell 174, 102–116 e114.

Ji, Y., Little, A.J., Banerjee, J.K., Hao, B., Oltz, E.M., Krangel, M.S., and Schatz, D.G. (2010). Promoters, enhancers, and transcription target RAG1 binding during V(D)J recombination. J Exp Med 207, 2809–2816.

Jung, D., Bassing, C.H., Fugmann, S.D., Cheng, H.L., Schatz, D.G., and Alt, F.W. (2003). Extrachromosomal recombination substrates recapitulate beyond 12/23 restricted VDJ recombination in nonlymphoid cells. Immunity 18, 65–74.

Khamlichi, A.A., and Feil, R. (2018). Parallels between Mammalian Mechanisms of Monoallelic Gene Expression. Trends Genet 34, 954–971.

Khor, B., and Sleckman, B.P. (2005). Intra-and inter-allelic ordering of T cell receptor beta chain gene assembly. Eur J Immunol 35, 964–970.

Kim, M.S., Chuenchor, W., Chen, X., Cui, Y., Zhang, X., Zhou, Z.H., Gellert, M., and Yang, W. (2018). Cracking the DNA Code for V(D)J Recombination. Mol Cell 70, 358–370 e354.

Kondilis-Mangum, H.D., Cobb, R.M., Osipovich, O., Srivatsan, S., Oltz, E.M., and Krangel, M.S. (2010). Transcription-dependent mobilization of nucleosomes at accessible TCR gene segments in vivo. J Immunol 184, 6970–6977.

Koralov, S.B., Novobrantseva, T.I., Konigsmann, J., Ehlich, A., and Rajewsky, K. (2006). Antibody repertoires generated by VH replacement and direct VH to JH joining. Immunity 25, 43–53.

Kwon, J., Imbalzano, A.N., Matthews, A., and Oettinger, M.A. (1998). Accessibility of nucleosomal DNA to V(D)J cleavage is modulated by RSS positioning and HMG1. Mol Cell 2, 829–839.

Larijani, M., Yu, C.C., Golub, R., Lam, Q.L., and Wu, G.E. (1999). The role of components of recombination signal sequences in immunoglobulin gene segment usage: a V81x model. Nucleic Acids Res 27, 2304–2309.

Lee, K.D., and Bassing, C.H. (2020). Two Successive Inversional Vbeta Rearrangements on a Single Tcrb Allele Can Contribute to the TCRbeta Repertoire. J Immunol 204, 78–86.

Levin-Klein, R., and Bergman, Y. (2014). Epigenetic regulation of monoallelic rearrangement (allelic exclusion) of antigen receptor genes. Front Immunol 5, 625.

Liang, H.E., Hsu, L.Y., Cado, D., Cowell, L.G., Kelsoe, G., and Schlissel, M.S. (2002). The “dispensable” portion of RAG2 is necessary for efficient V-to-DJ rearrangement during B and T cell development. Immunity 17, 639–651.

Lin, W.C., and Desiderio, S. (1994). Cell cycle regulation of V(D)J recombination-activating protein RAG-2. Proc Natl Acad Sci U S A 91, 2733–2737.

Livak, F., Burtrum, D.B., Rowen, L., Schatz, D.G., and Petrie, H.T. (2000). Genetic modulation of T cell receptor gene segment usage during somatic recombination. J Exp Med 192, 1191–1196.

Lovely, G.A., Brewster, R.C., Schatz, D.G., Baltimore, D., and Phillips, R. (2015). Single-molecule analysis of RAG-mediated V(D)J DNA cleavage. Proc Natl Acad Sci U S A 112, E1715–1723.

Majumder, K., Koues, O.I., Chan, E.A., Kyle, K.E., Horowitz, J.E., Yang-Iott, K., Bassing, C.H., Taniuchi, I., Krangel, M.S., and Oltz, E.M. (2015). Lineage-specific compaction of Tcrb requires a chromatin barrier to protect the function of a long-range tethering element. J Exp Med 212, 107–120.

Malissen, M., McCoy, C., Blanc, D., Trucy, J., Devaux, C., Schmitt-Verhulst, A.M., Fitch, F., Hood, L., and Malissen, B. (1986). Direct evidence for chromosomal inversion during T-cell receptor beta-gene rearrangements. Nature 319, 28–33.

Mombaerts, P., Clarke, A.R., Rudnicki, M.A., Iacomini, J., Itohara, S., Lafaille, J.J., Wang, L., Ichikawa, Y., Jaenisch, R., Hooper, M.L., et al. (1992). Mutations in T-cell antigen receptor genes alpha and beta block thymocyte development at different stages. Nature 360, 225–231.

Mostoslavsky, R., Alt, F.W., and Rajewsky, K. (2004). The lingering enigma of the allelic exclusion mechanism. Cell 118, 539–544.

Mostoslavsky, R., Singh, N., Tenzen, T., Goldmit, M., Gabay, C., Elizur, S., Qi, P., Reubinoff, B.E., Chess, A., Cedar, H., et al. (2001). Asynchronous replication and allelic exclusion in the immune system. Nature 414, 221–225.

Nadel, B., Tang, A., Escuro, G., Lugo, G., and Feeney, A.J. (1998). Sequence of the spacer in the recombination signal sequence affects V(D)J rearrangement frequency and correlates with nonrandom Vkappa usage in vivo. J Exp Med 187, 1495–1503.

Olaru, A., Patterson, D.N., Cai, H., and Livak, F. (2004). Recombination signal sequence variations and the mechanism of patterned T-cell receptor-beta locus rearrangement. Mol Immunol 40, 1189–1201.

Outters, P., Jaeger, S., Zaarour, N., and Ferrier, P. (2015). Long-Range Control of V(D)J Recombination & Allelic Exclusion: Modeling Views. Adv Immunol 128, 363–413.

Padovan, E., Casorati, G., Dellabona, P., Meyer, S., Brockhaus, M., and Lanzavecchia, A. (1993). Expression of two T cell receptor alpha chains: dual receptor T cells. Science 262, 422–424.

Pernis, B., Chiappino, G., Kelus, A.S., and Gell, P.G. (1965). Cellular localization of immunoglobulins with different allotypic specificities in rabbit lymphoid tissues. J Exp Med 122, 853–876.

Powers, S.E., Mandal, M., Matsuda, S., Miletic, A.V., Cato, M.H., Tanaka, A., Rickert, R.C., Koyasu, S., and Clark, M.R. (2012). Subnuclear cyclin D3 compartments and the coordinated regulation of proliferation and immunoglobulin variable gene repression. J Exp Med 209, 2199–2213.

Ramsden, D.A., and Wu, G.E. (1991). Mouse kappa light-chain recombination signal sequences mediate recombination more frequently than do those of lambda light chain. Proc Natl Acad Sci U S A 88, 10721–10725.

Ru, H., Chambers, M.G., Fu, T.M., Tong, A.B., Liao, M., and Wu, H. (2015). Molecular Mechanism of V(D)J Recombination from Synaptic RAG1-RAG2 Complex Structures. Cell 163, 1138–1152.

Ryu, C.J., Haines, B.B., Lee, H.R., Kang, Y.H., Draganov, D.D., Lee, M., Whitehurst, C.E., Hong, H.J., and Chen, J. (2004). The T-cell receptor beta variable gene promoter is required for efficient V beta rearrangement but not allelic exclusion. Mol Cell Biol 24, 7015–7023.

Sarukhan, A., Garcia, C., Lanoue, A., and von Boehmer, H. (1998). Allelic inclusion of T cell receptor alpha genes poses an autoimmune hazard due to low-level expression of autospecific receptors. Immunity 8, 563–570.

Schatz, D.G., and Swanson, P.C. (2011). V(D)J recombination: mechanisms of initiation. Annu Rev Genet 45, 167–202.

Schlimgen, R.J., Reddy, K.L., Singh, H., and Krangel, M.S. (2008). Initiation of allelic exclusion by stochastic interaction of Tcrb alleles with repressive nuclear compartments. Nat Immunol 9, 802–809.

Schuldt, N.J., Auger, J.L., Spanier, J.A., Martinov, T., Breed, E.R., Fife, B.T., Hogquist, K.A., and Binstadt, B.A. (2017). Cutting Edge: Dual TCRalpha Expression Poses an Autoimmune Hazard by Limiting Regulatory T Cell Generation. J Immunol 199, 33–38.

Serwold, T., Hochedlinger, K., Inlay, M.A., Jaenisch, R., and Weissman, I.L. (2007). Early TCR expression and aberrant T cell development in mice with endogenous prerearranged T cell receptor genes. J Immunol 179, 928–938.

Shih, H.Y., and Krangel, M.S. (2013). Chromatin architecture, CCCTC-binding factor, and V(D)J recombination: managing long-distance relationships at antigen receptor loci. J Immunol 190, 4915–4921.

Shinkai, Y., Rathbun, G., Lam, K.P., Oltz, E.M., Stewart, V., Mendelsohn, M., Charron, J., Datta, M., Young, F., Stall, A.M., et al. (1992). RAG-2-deficient mice lack mature lymphocytes owing to inability to initiate V(D)J rearrangement. Cell 68, 855–867.

Sicinska, E., Aifantis, I., Le Cam, L., Swat, W., Borowski, C., Yu, Q., Ferrando, A.A., Levin, S.D., Geng, Y., von Boehmer, H., et al. (2003). Requirement for cyclin D3 in lymphocyte development and T cell leukemias. Cancer Cell 4, 451–461.

Skok, J.A., Gisler, R., Novatchkova, M., Farmer, D., de Laat, W., and Busslinger, M. (2007). Reversible contraction by looping of the Tcra and Tcrb loci in rearranging thymocytes. Nat Immunol 8, 378–387.

Sleckman, B.P., Bassing, C.H., Hughes, M.M., Okada, A., D’Auteuil, M., Wehrly, T.D., Woodman, B.B., Davidson, L., Chen, J., and Alt, F.W. (2000). Mechanisms that direct ordered assembly of T cell receptor beta locus V, D, and J gene segments. Proc Natl Acad Sci U S A 97, 7975–7980.

Steinel, N.C., Brady, B.L., Carpenter, A.C., Yang-Iott, K.S., and Bassing, C.H. (2010). Posttranscriptional silencing of VbetaDJbetaCbeta genes contributes to TCRbeta allelic exclusion in mammalian lymphocytes. J Immunol 185, 1055–1062.

Steinel, N.C., Fisher, M.R., Yang-Iott, K.S., and Bassing, C.H. (2014). The ataxia telangiectasia mutated and cyclin D3 proteins cooperate to help enforce TCRbeta and IgH allelic exclusion. J Immunol 193, 2881–2890.

Steinel, N.C., Lee, B.S., Tubbs, A.T., Bednarski, J.J., Schulte, E., Yang-Iott, K.S., Schatz, D.G., Sleckman, B.P., and Bassing, C.H. (2013). The ataxia telangiectasia mutated kinase controls Igkappa allelic exclusion by inhibiting secondary Vkappa-to-Jkappa rearrangements. J Exp Med 210, 233–239.

Tillman, R.E., Wooley, A.L., Khor, B., Wehrly, T.D., Little, C.A., and Sleckman, B.P. (2003). Cutting edge: targeting of V beta to D beta rearrangement by RSSs can be mediated by the V(D)J recombinase in the absence of additional lymphoid-specific factors. J Immunol 170, 5–9.

VanDyk, L.F., Wise, T.W., Moore, B.B., and Meek, K. (1996). Immunoglobulin D(H) recombination signal sequence targeting: effect of D(H) coding and flanking regions and recombination partner. J Immunol 157, 4005–4015.

Vettermann, C., and Schlissel, M.S. (2010). Allelic exclusion of immunoglobulin genes: models and mechanisms. Immunol Rev 237, 22–42.

von Boehmer, H., and Melchers, F. (2010). Checkpoints in lymphocyte development and autoimmune disease. Nat Immunol 11, 14–20.

Wang, X., Xiao, G., Zhang, Y., Wen, X., Gao, X., Okada, S., and Liu, X. (2008). Regulation of Tcrb recombination ordering by c-Fos-dependent RAG deposition. Nat Immunol 9, 794–801.

Wei, Z., and Lieber, M.R. (1993). Lymphoid V(D)J recombination. Functional analysis of the spacer sequence within the recombination signal. J Biol Chem 268, 3180–3183.

Wilson, A., Marechal, C., and MacDonald, H.R. (2001). Biased V beta usage in immature thymocytes is independent of DJ beta proximity and pT alpha pairing. J Immunol 166, 51–57.

Wu, C., Bassing, C.H., Jung, D., Woodman, B.B., Foy, D., and Alt, F.W. (2003). Dramatically increased rearrangement and peripheral representation of Vbeta14 driven by the 3’Dbeta1 recombination signal sequence. Immunity 18, 75–85.

Wu, C., Ranganath, S., Gleason, M., Woodman, B.B., Borjeson, T.M., Alt, F.W., and Bassing, C.H. (2007). Restriction of endogenous T cell antigen receptor beta rearrangements to Vbeta14 through selective recombination signal sequence modifications. Proc Natl Acad Sci U S A 104, 4002–4007.

Yang-Iott, K.S., Carpenter, A.C., Rowh, M.A., Steinel, N., Brady, B.L., Hochedlinger, K., Jaenisch, R., and Bassing, C.H. (2010). TCR beta feedback signals inhibit the coupling of recombinationally accessible V beta 14 segments with DJ beta complexes. J Immunol 184, 1369–1378.

Zal, T., Weiss, S., Mellor, A., and Stockinger, B. (1996). Expression of a second receptor rescues self-specific T cells from thymic deletion and allows activation of autoreactive effector function. Proc Natl Acad Sci U S A 93, 9102–9107.

