## Supplemental Figures for "Poor Quality Vβ Recombination Signal Sequences Enforce TCRβ Allelic Exclusion by Limiting the Frequency of Vβ Recombination"

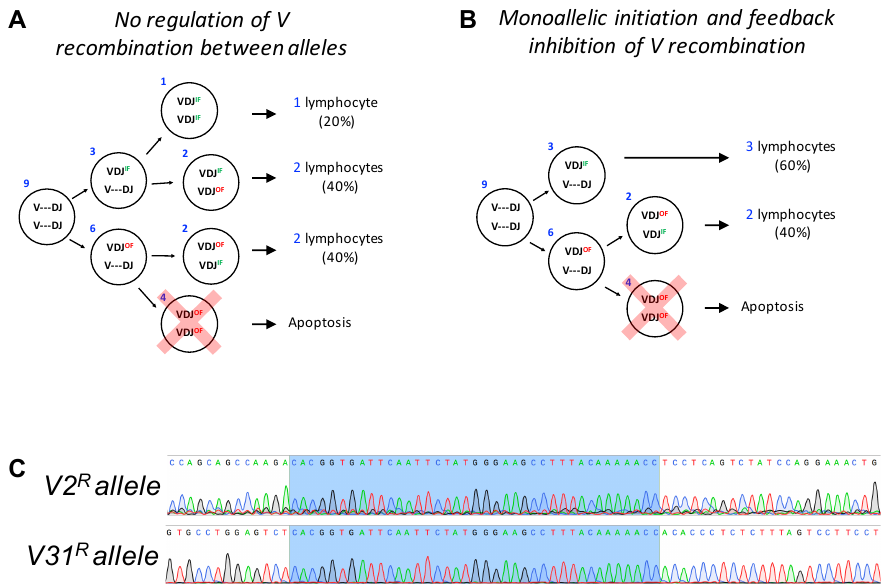

**Figure S1. Models of V recombination and validation of *V2* and *V31* RSS replacement mice.**

(A and B) Theoretical frequencies of V(D)J rearrangements and their reading frame status in lymphocytes assuming no regulation of V recombination between alleles (A) or mono-allelic initiation and feedback inhibition of V recombination (B).

(C) Sequence validation of the V2 or V31 RSS replacement with the 3’Dβ1 RSS. The 3’Dβ1 RSSs are highlighted in blue.

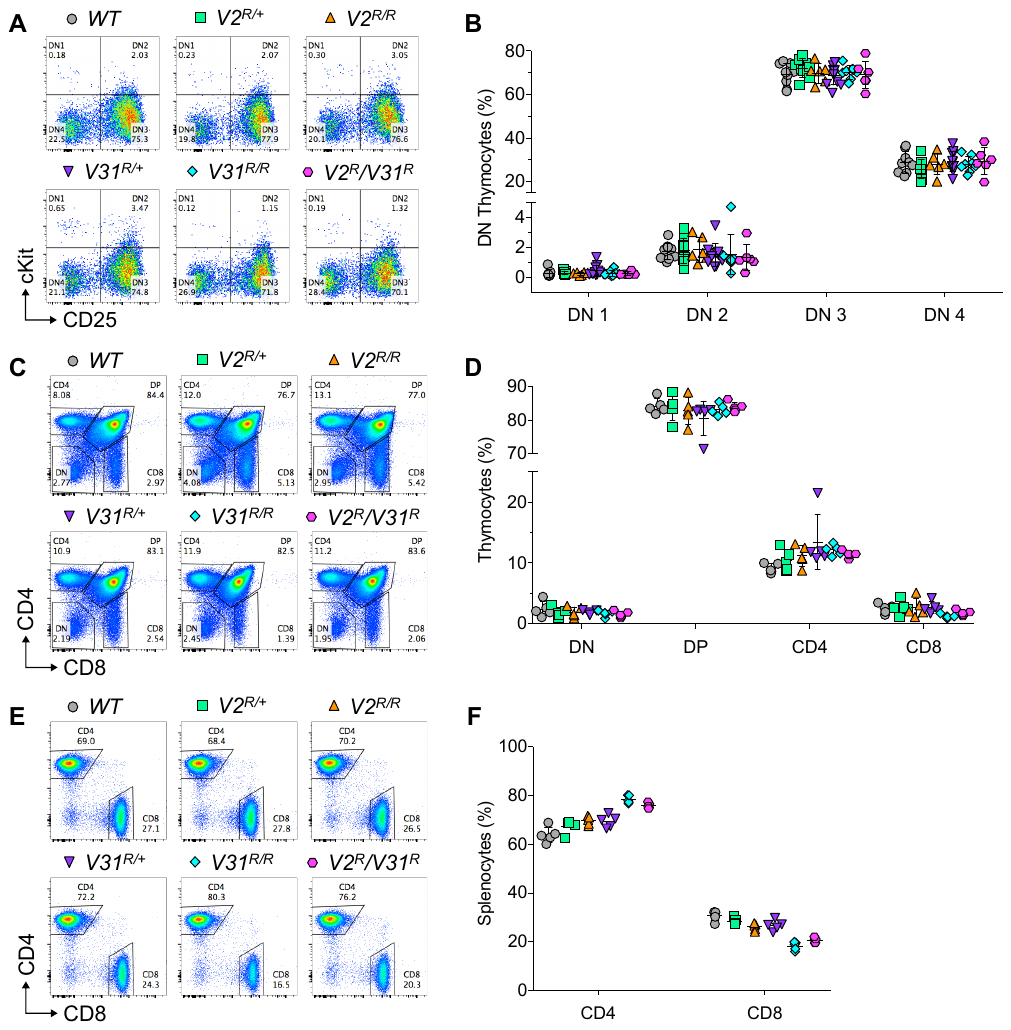

**Figure S2. Normal Gross αβ T cell development in *V2* and *V31* RSS replacement mice.**

(A and B) DN thymocyte development. Representative plots (A) and quantification (B) of DN cells. Gated on Lin^-^CD4^-^CD8^-^TCRβ^-^ thymocytes (n ≥ 5).

(C and D) Global thymocyte development. Representative plots (C) and quantification (D) of DN, DP, CD4^+^, and CD8^+^ thymocytes (n = 5).

(E and F) Representative plots (E) and quantification (F) of SP αβ T cells in the spleen. Gated on TCRβ^+^ cells (n = 5).

All quantification plots show mean ± SD.

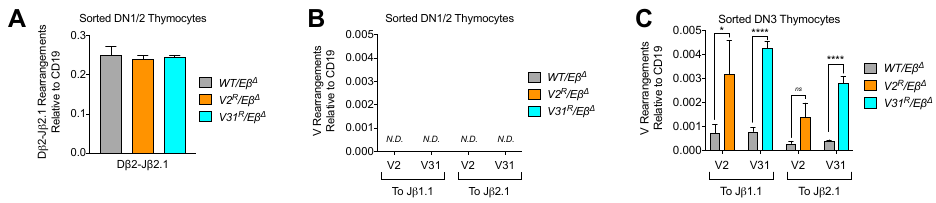

**Figure S3. The 3’Dβ1 RSS replacement increases *V2* and *V31* recombination in DN3 thymocytes.**

(A) Quantification of Dβ2-Jβ2.1 rearrangements by TaqMan qPCR in DN1/2 thymocytes (n = 3).

(B and C) Quantification of indicated Vβ rearrangements by TaqMan qPCR in DN1/2 (B) or DN3 (C) thymocytes (n = 3, two-way ANOVA).

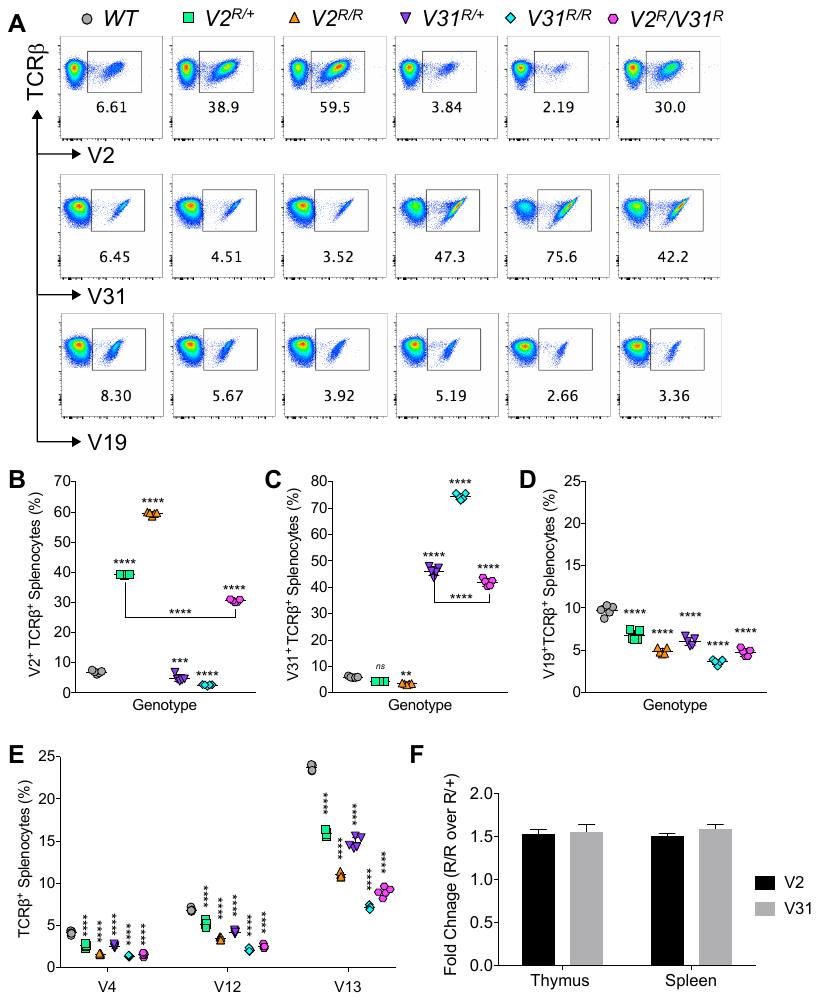

**Figure S4.** **Peripheral αβ T cells exhibit similar shifts in the Vβ repertoire in RSS replacement mice.**

(A) Representative plots of SP splenocytes expressing V2^+^, V31^+^, or V19^+^ TCRβ chains.

(B-D) Quantification of V2^+^ (B), V31^+^ (C), or V19^+^ (D) SP thymocytes (n = 5, one-way ANOVA).

(E) Quantification of SP splenocytes expressing V4^+^, V12^+^, or V13^+^ TCRβ chains (n = 5, two-way ANOVA).

(F) Ratio of the V2^+^ and V31^+^ Vβ repertoires. The fold change calculates the frequency of V2^+^ cells from *V2^R/R^* mice divided by *V2^R/+^* mice. A similar calculation was made for V31^+^ cells from *V31^R/R^* and *V31^R/+^* mice.

All quantification plots show mean ± SD. Multiple post-tests are compared to *WT* unless indicated by bars, and *p*-values are corrected for multiple tests. ns=not significant, ***p<0.01*, ****p<0.001*, *****p<0.0001*.

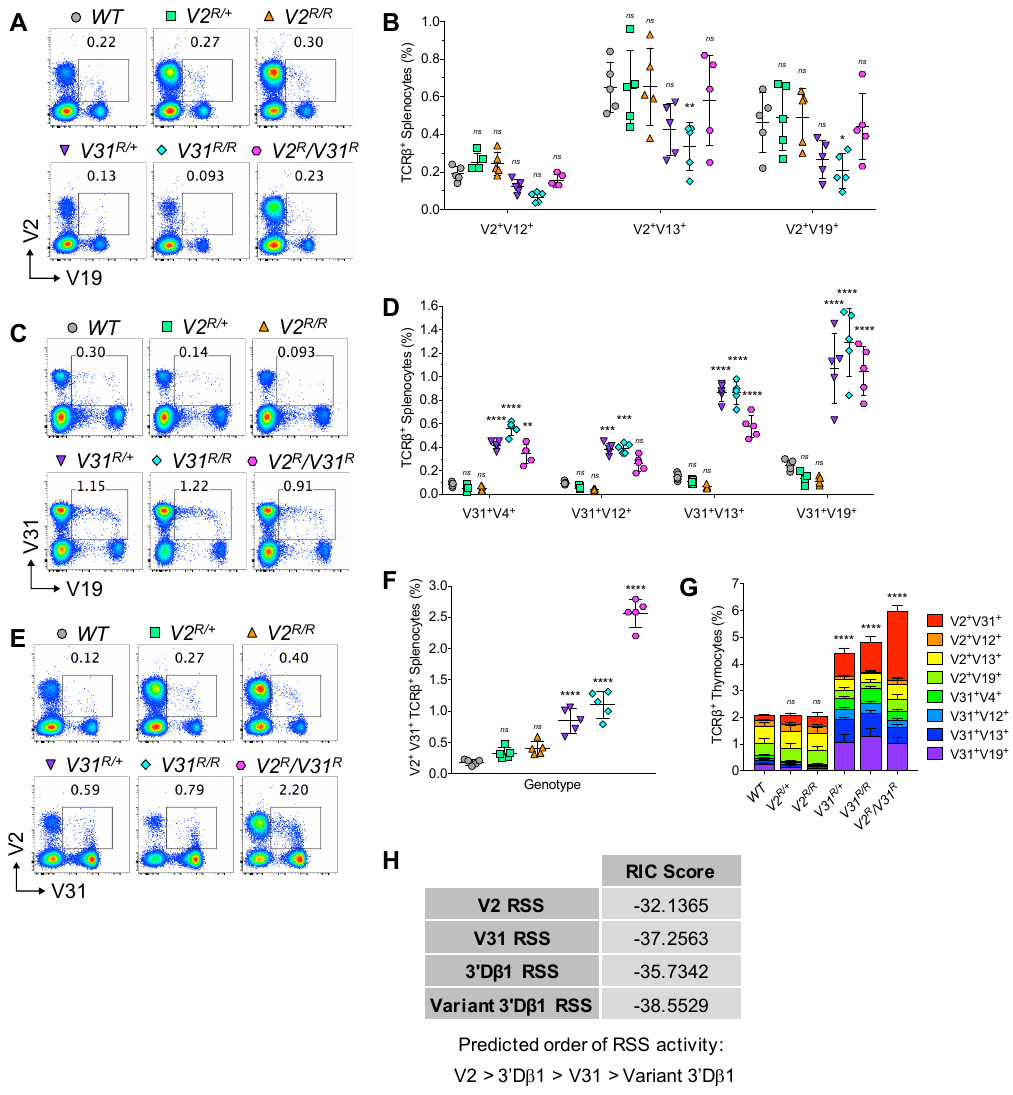

**Figure S5. αβ T cells exhibiting biallelic *Tcrb* gene expression seed the periphery.**

(A, C, and E) Representative plots of SP splenocytes expressing both V2^+^ and V19^+^ (A), V31^+^ and V19^+^ (B), or V2^+^ and V31^+^ (E) TCRβ chains.

(B, D, and F) Quantification of SP splenocytes expressing the two indicated TCRβ chains. [n = 5, two-way ANOVA for (B) and (D), one-way ANOVA for (F)].

(G) Quantification of double-staining SP splenocytes for each Vβ combination tested (n = 5, two-way ANOVA).

(H) Recombination information content (RIC) scores of RSSs in this study.

All quantification plots show mean ± SD. Multiple post-tests are compared to *WT* unless indicated by bars, and *p*-values are corrected for multiple tests. ns=not significant, **p<0.05*, ***p<0.01*, ****p<0.001*, *****p<0.0001*.

**Supplementary Table 1. Analysis of *Tcrb* rearrangements in *Tcrb^Tg^* T cell hybridomas**

| Genotype and Source | *Tcrb^Tg^** | | *V2^R/+^* | | *Tcrb^Tg^V2^R^* | | *V31^R/+^* | | *Tcrb^Tg^V31^R^* | |
| --- | --- | --- | --- | --- | --- | --- | --- | --- | --- | --- |
|  | Number | % | Number | % | Number | % | Number | % | Number | % |
| Clonal Hybridomas Assayed | 129 | 100.0 | 18 | 100.0 | 56 | 100.0 | 133 | 100.0 | 188 | 100.0 |
| V(D)J | 12 | 9.3 |  |  |  |  |  |  | 28 | 14.9 |
| V2(D)J |  |  | 9 | 50.0 | 0 | 0.0 |  |  |  |  |
| V31(D)J | 3 | 2.3 |  |  |  |  | 66 | 49.6 | 28 | 14.9 |
| V31-to-DJ | 3 | 2.3 |  |  |  |  |  |  | 24 | 12.8 |
| V31-to-J |  |  |  |  |  |  |  |  | 4 | 2.1 |
| V31(D)J on one allele |  |  |  |  |  |  | 35 | 100.0 |  |  |
| V31-to-DJ |  |  |  |  |  |  | 25 | 71.4 |  |  |
| V31-to-J |  |  |  |  |  |  | 10 | 28.6 |  |  |

* = Brady et al. 2010
